# Cross-sectional and longitudinal changes in category-selectivity in visual cortex following pediatric cortical resection

**DOI:** 10.1101/2024.12.08.627367

**Authors:** Tina T. Liu, Michael C. Granovetter, Anne Margarette S. Maallo, Sophia Robert, Jason Z. Fu, Christina Patterson, David C. Plaut, Marlene Behrmann

**Author notes:** Corresponding author: Marlene Behrmann. Joint first authorship (Tina T. Liu, Michael C. Granovetter).

## Abstract

The topographic organization of category-selective responses in human ventral occipitotemporal cortex (VOTC) and its relationship to regions subserving language functions is remarkably uniform across individuals. This arrangement is thought to result from the clustering of neurons responding to similar inputs, constrained by intrinsic architecture and tuned by experience. We examined the malleability of this organization in individuals with unilateral resection of VOTC during childhood for the management of drug-resistant epilepsy. In cross-sectional and longitudinal functional imaging studies, we compared the topography and neural representations of 17 category-selective regions in individuals with a VOTC resection, a ‘control patient’ with resection outside VOTC, and typically developing matched controls. We demonstrated both adherence to and deviation from the standard topography and uncovered fine-grained competitive dynamics between word- and face-selectivity over time in the single, preserved VOTC. The findings elucidate the nature and extent of cortical plasticity and highlight the potential for remodeling of extrastriate architecture and function.

**Teaser:** After pediatric cortical resection, deviations from the constraints of standard topography in visual cortex reflect plasticity.

## Introduction

The human visual system exhibits a topographic organization that is largely replicable and uniform across individuals and across languages and cultures (1). While primary visual cortex is homologous across the two cerebral hemispheres, each with low-level information of the contralateral visual field, ventral occipitotemporal cortex (VOTC) exhibits distinct patterns of functional selectivity for different categories of complex stimuli (e.g., faces, objects, words, scenes) both within and between hemispheres (2–4). This extrastriate topography is thought to reflect the clustering of neurons responding to functionally similar inputs, constrained by the intrinsic architecture of visual cortex (5–7), even in the absence of category-specific learning pressures (8). Efforts to elucidate the phylogenetic and ontogenetic origins of category-selective organization are ongoing, and fine-grained topographic maps in humans (4, 9) and in non-human primates (10) have already been identified (for recent review, see Bourne et al. (11)).

Notwithstanding the consistent and reliable characterization afforded by these topographic maps and their stereotypical relationships with other cortical areas, such as those subserving language function and regions of early visual cortex, the potential extent and nature of their plasticity remains to be determined. Beyond the maps and the spatial relations between the demarcated regions, it is also important to understand what information is instantiated in these regions and whether, for example, representational content is necessarily tied to a stereotypical location or is maintained even when the topography progressively deviates from the typical arrangement (e.g., after neural injury).

### The developmental emergence and uniformity of category selectivity

In humans, category-selective responses beyond early visual cortex emerge over development, with dorsal, parietal regions emerging and maturing seemingly earlier than ventral, temporal regions (12, 13). However, even within VOTC itself, some regions have an earlier sensitive period and evince category selectivity ahead of other regions (10, 11, 14). For example, bilateral object- and scene-selective regions appear to mature earlier (15, 16) than face- and word-selective regions (17), with these latter regions more critically dependent on visual experience (18, 19).

Indeed, word- and face-selective areas evolve over a protracted developmental trajectory, stabilizing by adulthood with a weighted asymmetry: words are largely represented just in left VOTC, while faces are more bilaterally represented, with stronger activation in the right VOTC (20) than in the left VOTC (21). One explanation for this prolonged trajectory is based on the high perceptual confusability between individual exemplars within the category of words (e.g., two similar words such as ‘hair’ and ‘lair’) and of faces (e.g., two similar faces such as those of Elvis Presley and George Clooney), which is less the case for other visual categories (e.g., objects, scenes). This prolonged acquisition of detailed representations for words and faces offers a special opportunity for quantifying plasticity over months and years by concurrent tracking of neural alterations and associated behavioral changes.

The pre-eminent word-selective area, the ‘Visual Word Form Area’ (VWFA), emerges in concert with literacy acquisition around ages five or six years (22) and is almost ubiquitously lateralized to the left hemisphere (LH) (1, 23), potentially via pressure to be spatially co-localized with LH-dominant language areas (24, 25). By contrast, the trajectory of the pre-eminent face- selective region, the ‘Fusiform Face Area’ (FFA), begins early in life (26, 27) and continues to be refined through early adulthood (28, 29). The FFA ultimately lateralizes predominantly to the right hemisphere (RH), either as a result of competition with the LH-lateralized VWFA once literacy is acquired (2, 30, 31) and/or via pressure to coordinate with other relevant RH-lateralized processes, including social processing (32). A recent study calculated the distribution of lateralization of word and face selectivity in the RH and/or LH from fMRI scans of 54 right- handed college-age individuals. Bilateral and LH word selectivity was observed in 5 and 47 individuals, respectively, and no individual showed only RH selectivity. Bilateral and RH face selectivity was noted in 19 and 34 individuals, respectively, and no individual showed only LH selectivity. Last, for common objects, bilateral, unilateral RH, and unilateral LH selectivity was observed in 42, 10, and 2 individuals, respectively. Altogether, the findings demonstrated that by early adulthood, most individuals show a LH bias for words and a RH bias for faces, and that the hemispheric specialization is specific to words and faces (32).

The marked LH lateralization and prolonged emergence of word representations offer a unique opportunity to study the principles that govern the ontogenesis of the VWFA, especially because word selectivity is too recent evolutionarily to be innately predetermined (1, 23) and, thus, is unlikely to be specified in the genome (33). That a word-selective cortical region can be reliably identified in the LH of humans already attests to the malleability of human cortex, as does the fact that left-handed individuals, especially those with RH-language dominance, do not show the typical LH-lateralization of VWFA (34, 35). However, the fact that the VWFA is so replicable across the population at large (1) and is independent of the native tongue of the reader (36), raises many questions concerning the constraints governing this relatively new cultural ability.

### The current study

The primary question addressed here concerns the emergence of category-selective regions in VOTC, including those associated with word and face processing, the malleability of their hemispheric lateralization, localization within the hemisphere, their relationship to other key areas (e.g., those subserving language function and early visual cortex), and their representational specificity. To address this question, we leverage data from a unique participant population that allows us to examine the plasticity of category-selective VOTC, both cross-sectionally and longitudinally, and across both the LH and RH. This population is comprised of individuals who have undergone a unilateral childhood resection of VOTC for the management of drug-resistant epilepsy (DRE), presumably resulting in pressure on the preserved cortex to accommodate functions of the resected tissue. Because plasticity is thought to be greater in children than in adults (37–40), the study of these individuals affords a distinct opportunity to monitor the evolving category topography. Moreover, there are many questions but few studies determining whether the represented category-selective information is necessarily contingent on the topographic site of the category. For example, if the VWFA emerges in the RH of an individual with a LH resection, do the neural representations within this atypical RH VWFA correspond to those of the standard LH-lateralized VWFA? On some accounts, the presence of a category- selective region need not precede the evolution of more refined neural representations (41), and distributed representational patterns may even scaffold the later emergence of univariate category selectivity (42).

In our previous research with individuals from this rather rare population, we showed that a group of 39 patients with complete hemispheric surgery during childhood scored, on average, 85% accuracy in word and in face discrimination which, although statistically inferior to the 92% accuracy of the typically developing (TD) controls, was better than predicted based on the extent of the anatomical resection (43) (see also Simmons et al. (44)). Using fMRI, we identified a few category-selective regions in the preserved VOTC of children with unilateral VOTC resection and found that their category-selective representations were largely similar to those control patients with a resection outside VOTC and to matched typical controls (45). Also, in a limited longitudinal case study of a patient with RH VOTC resection, category-selective development mirrored that of cross-sectional controls, except in the left FFA (46). Last, we observed normal fMRI repetition suppression for faces, words, and objects in patients’ single VOTC but quantitatively poorer behavioral accuracy scores than typical controls, suggesting that a single hemisphere alone does not suffice for normal visual recognition behavior despite intact unilateral neural signatures for visual exemplar individuation (47).

Our prior work was restricted to a limited number of category-selective regions and did not examine alterations in VWFA lateralization in relation to language regions. Most importantly, as word representations typically emerge over development in left VOTC, a critical desideratum is whether one can observe the microgenesis of competition between word and face representations evolve over time in the RH after left VOTC resection. Capturing longitudinal changes in right VOTC under the extreme constraint of developing without a left VOTC would attest to the inherent plasticity of VOTC. Thus, here, we build on these foundations and compare the widespread spatial topography of 17 category-selective regions, their hemispheric lateralization, relationship to lateralized language regions and early visual cortex, and their representational bases in five individuals with a VOTC childhood resection (see Figure 1). Three individuals underwent resections that encompassed the left posterior VOTC (KN, SN, and TC; Figure 1A), one has a right posterior VOTC resection (UD; Figure 1B), and one ’control patient’, OT, has a left anterior temporal resection (e.g., outside VOTC; Figure 1C). Additionally, in three of the patients (TC, UD, and OT), we characterize the longitudinal neural profile over multiple fMRI sessions, and, for TC, the longitudinal data span pre- to post-surgery. Triangulating multiple dimensions of the neural profile as a function of resection site (left versus right and anterior vs. posterior) using both cross-sectional and longitudinal approaches offers important insights into the malleability of VOTC’s organization and the dynamics by which plasticity may operate.

**Figure 1.**
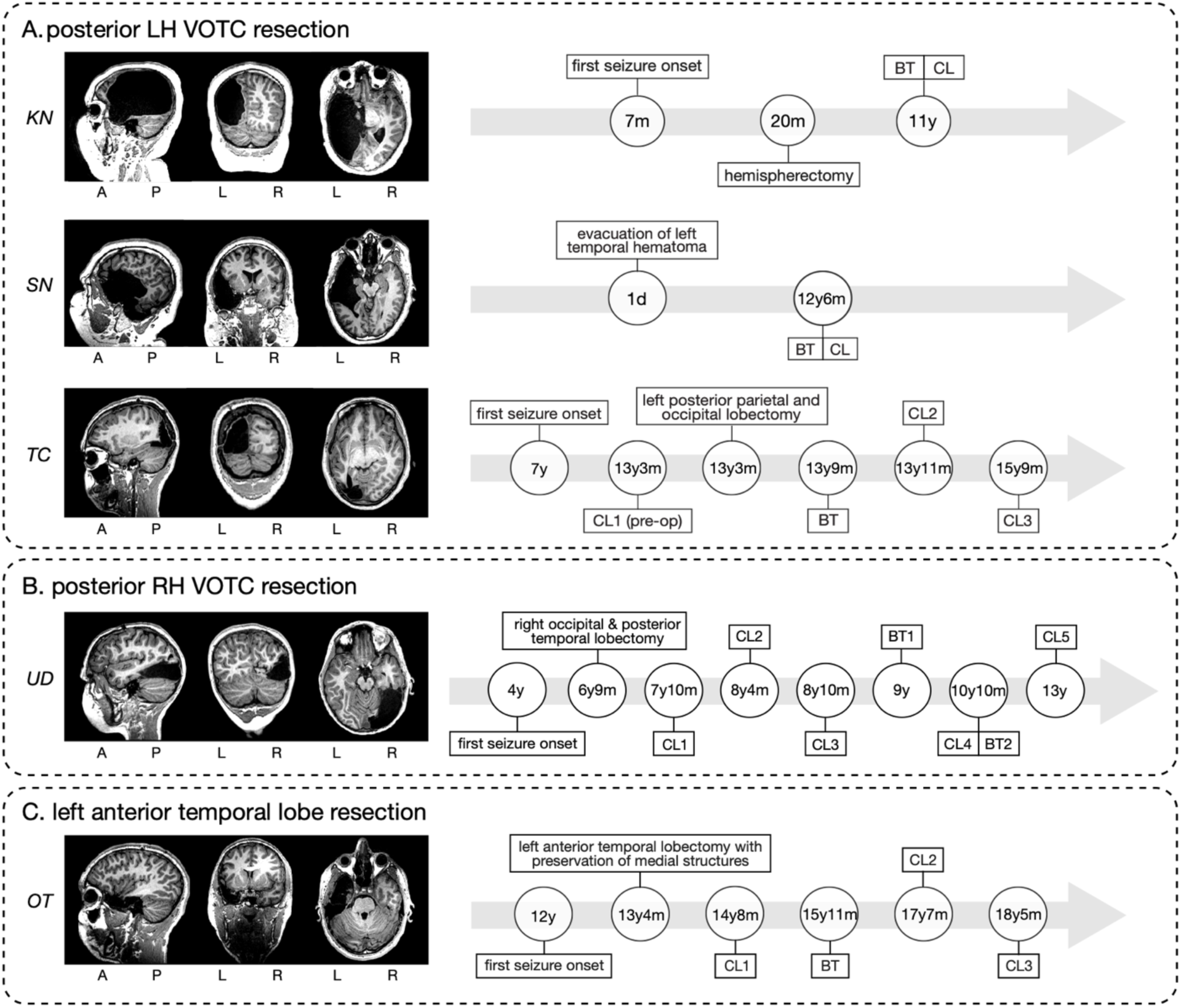
Postoperative structural MRI scans for the five pediatric resection patients in our study (left) and, for each, the time course of the behavioral testing (BT) and functional imaging sessions using a category localizer (CL) (right). A = anterior, P = posterior, L = left hemisphere, R = right hemisphere. d, m, y = days, months, years of age. **(A)** posterior LH VOTC resection patients: KN, SN, and TC. **(B)** posterior RH VOTC resection patient: UD. **(C)** ’control’ (e.g., outside VOTC) left anterior temporal lobe resection patient: OT.

## Results

We first characterize the visual behavior of the patients and TD controls and then analyze the cross-sectional and longitudinal fMRI data of category-selective regions of interest (ROIs) in VOTC.

### Visual behavior performance

To evaluate perceptual competence, participants completed two intermediate-level and two high-level vision tasks. Table 1 reports patients’ scores and whether they show a deficit relative to the TD control distribution, as determined using two-tailed Crawford’s modified t tests for single subjects versus a group with p < .05 (48).

**Table 1.**
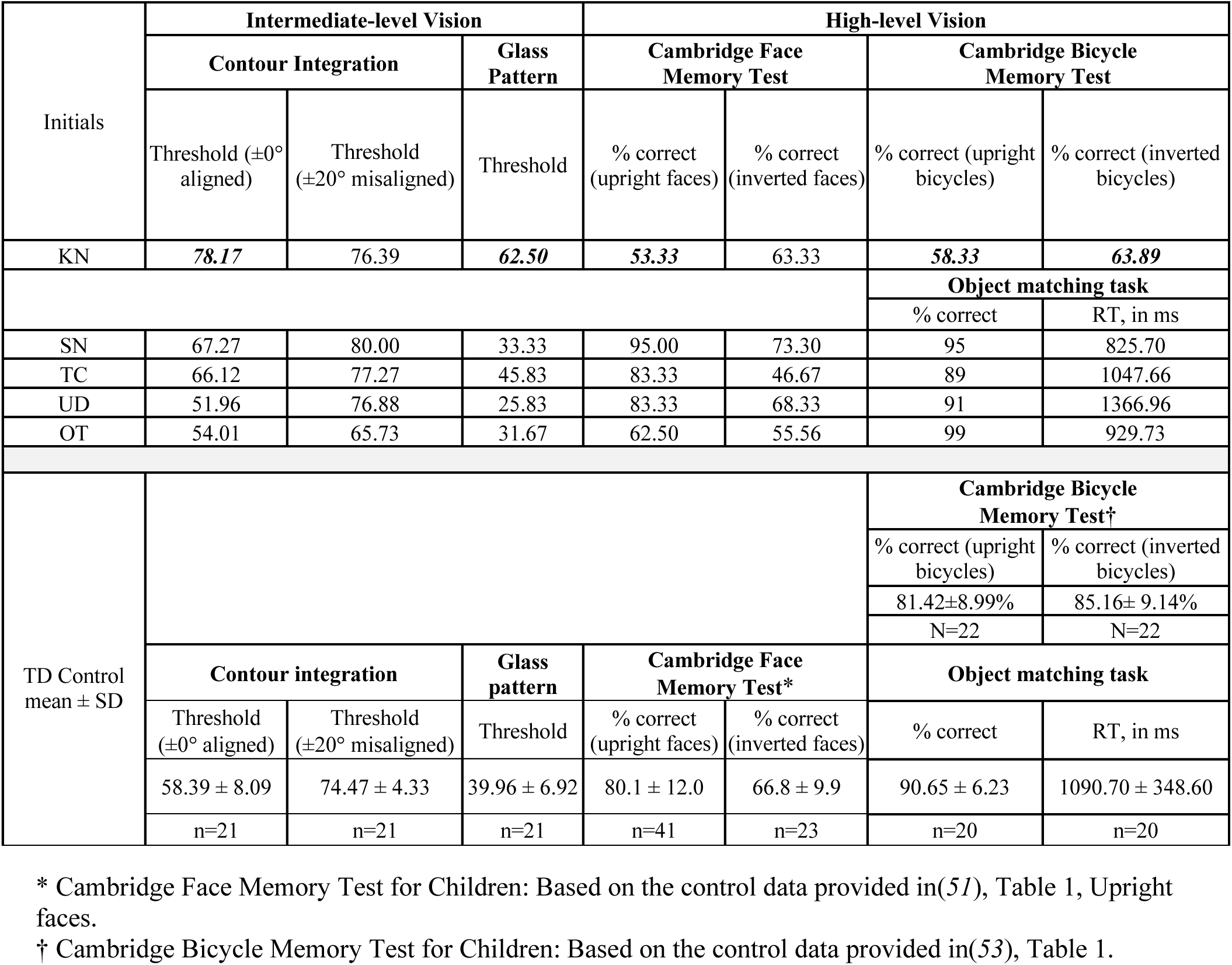
Results of visual perceptual behavior in five patients and the average performance in TD controls. Numbers in bold and italic font (only present in KN) denote significant deviations from the TD controls’ performance.

For assessing intermediate vision, we measured thresholds in a contour integration task in which we presented aligned or misaligned Gabor patches (in separate blocks), and participants located the ’egg’ shape in the display (49) (see Figure S1A). All patients’ thresholds fell within the TD control range except for KN in the aligned condition (Table 1). The same result held for thresholds for detecting which of consecutively presented glass pattern stimuli had more concentricity (50) (see Figure S1B and Table 1).

Accuracy was normal for all patients on high-level vision tasks except, again, for KN on the upright faces on the Cambridge Face Memory Test for Children (51) (see Figure S1C and Table 1). The four patients, SN, TC, UD, and OT, who completed the object matching task (52) (see Figure S1D) showed accuracy within normal limits (Table 1). KN, who completed the Cambridge Bicycle Memory Test for Children (53) (see Figure S1E), performed outside the TD range (Table 1). In summary, we observed typical intermediate- and high-level visual perception in all patients except for KN.

### fMRI of category-selective ROIs

Prior to analyzing the fMRI data, we determined that there were no significant differences between the data from patients versus controls in terms of head motion or average temporal signal-to-noise ratio (tSNR) across all functional voxels (see Methods for further details). This ensures the equivalence of data quality across the groups, and affirms that any observed group differences are unlikely to be due to the data acquisition process itself.

#### Category selectivity and topographic mapping

Using a functional category localizer to identify ROIs for assessing widespread VOTC topography (45, 46) (Figure 2A), we mapped 17 ROIs that are preferentially responsive to faces, scenes, objects, words, or scrambled objects in each of the 25 TD controls (Figures 2B-C and S3). The regions included the bilateral face-selective FFA (54, 55) and posterior superior temporal sulcus (56) (STS); bilateral scene-selective parahippocampal place area (57) (PPA) and transverse occipital sulcus (58) (TOS; also referred to as OPA); bilateral object-selective lateral occipital complex (59) (LOC) consisting of lateral occipital cortex (LO) and posterior fusiform (60, 61) (pF); left-lateralized word-selective VWFA (62), inferior frontal gyrus (IFG), and superior temporal gyrus (STG); and bilateral early visual cortex (EVC).

**Figure 2.**
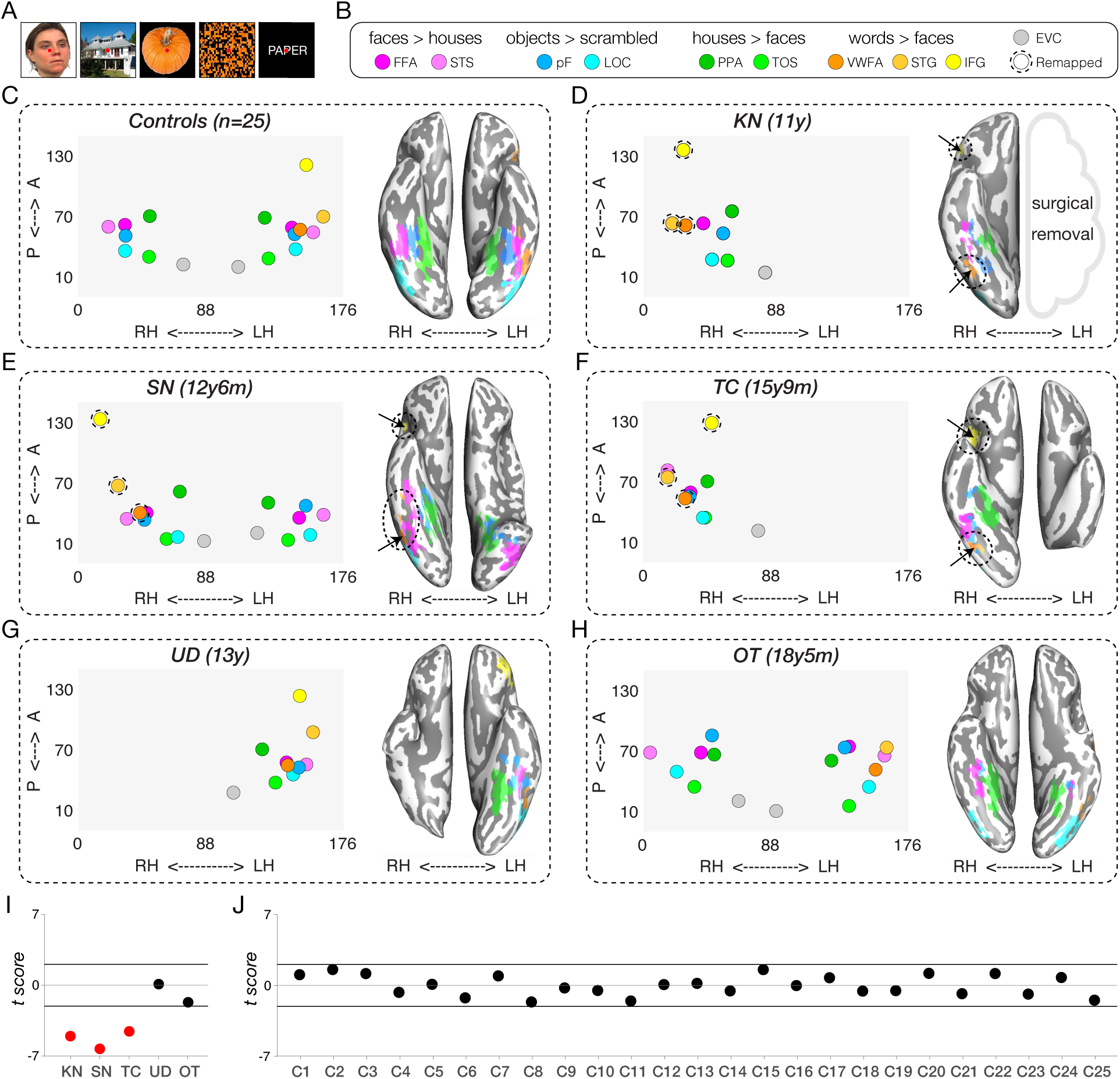
Spatial organization (in native space) of category selectivity in TD controls and patients. **(A)** Example stimuli used in the functional localizer experiment (see Methods for details). **(B)** Contrasts to define category-selective activations for each region. FFA = fusiform face area, STS = superior temporal sulcus, pF = posterior fusiform, LOC = lateral occipital complex, PPA = parahippocampal place area, TOS = transverse occipital sulcus, VWFA = visual word form area, STG = superior temporal gyrus, IFG = inferior frontal gyrus, EVC = early visual cortex. (**C-H**) Category-selective regions of interest (ROIs) in the control group (averaged across participants; n=25) and from the last scan session in each patient. The left side within each panel visualizes the average spatial distribution of category-selective ROIs in the controls and in each patient. The x-axis represents coordinates in the medial-lateral direction for each hemisphere (left: 88-176, right: 0-88 in native space), and the y-axis represents coordinates in the anterior-posterior direction. Filled colored circles indicate ROIs that can be identified in this scan; circles surrounded by dotted lines represent ROIs for word function that are typically left-lateralized but here are localized to the RH. The right side within each panel visualizes the ventral category-selective activations on the inflated cortical surface (ventral view) with corresponding dotted ovals indicating atypical sites of activation (word-selective ROIs in RH). See Figure S2 for details of the ROIs that are resected, not covered, or not found in the patients. See also Figure 4 for spatial organization of category selectivity in different scan sessions involving longitudinal patients TC, UD, and OT. A = anterior, P = posterior, LH = left hemisphere, RH = right hemisphere. Note that the left panel depicts both ventral and dorsal-lateral ROIs, but only the ventral ROIs are shown in the ventral view of the inflated surface in the right panel of B-H. **(I)** Crawford’s modified t score of difference in the spatial organization between each patient and the TD control group. **(J)** Crawford’s modified t score of difference in the spatial organization between each control and all other TD controls. See also Figure S3 for the spatial organization maps of each individual TD control.

In the patients, the number of identifiable category-selective ROIs varied, either because of resection or absence of functional activation (Figure S2). The coordinates of identified ROIs (posterior to anterior and left to right) in native volume space are shown in the left panel within Figure 2D-H for one scan session per patient (most recent if scanned longitudinally). In summary, for KN (left hemispherectomy), we identified all category-selective ROIs in the preserved RH except for STS (Figures 2D and S2), including the right-lateralized VWFA, STG, and IFG. In SN (left temporal resection), ROIs for all categories were present bilaterally, except for VWFA, STG, and IFG, which were all localized to the RH (Figures 2E and S2). In TC (left posterior occipitotemporal and parietal resection), we detected category-selective ROIs only in the RH, including VWFA, STG, and IFG (Figures 2F and S2). In UD (right VOTC resection), all category-selective ROIs were localized but only in the LH (Figures 2G and S2). Last, in control patient OT (left anterior temporal resection), all category-selective ROIs within VOTC were successfully identified bilaterally with the (standard) LH-lateralized VWFA and STG, except that the IFG was not covered in the limited brain coverage across three longitudinal scans (as we prioritized covering the anterior temporal lobe of OT’s intact hemisphere).

### Cross-sectional analysis

#### Spatial topography of category selectivity

Having determined which category-selective ROIs were identified in each participant, we then evaluated their spatial organization. In typical individuals, the EVC, PPA, pF, FFA, and VWFA are stereotypically organized along a medial-lateral axis within the ventral visual pathway in each hemisphere. To assess whether the patients’ ventral visual pathway obeys this medial-lateral bias (63, 64), we first extracted the native coordinates of the peak voxel within the ventral ROIs (EVC, PPA, pF, FFA, VWFA) for each of 25 age-matched fMRI controls (see Figure S3). Next, we computed the correlation between the x-coordinates of each patient’s available ventral ROIs to the corresponding average x-coordinates in the TD controls (Figure 2C). Crawford’s modified t tests, comparing individual patient’s correlation values to the respective TD control distribution, revealed significant deviations in all three LH VOTC resection cases (KN, SN, and TC: all |t(_24_)| values > 5.53; Figure 2I, red dots), resulting from the atypical presence of the VWFA, STG, and IFG in the RH (Figure 2D-F). In contrast, UD and OT showed typical medial-lateral organization of category selectivity in all scan sessions (Figure 2I, black dots). The same analysis for each individual control, compared against the other 24 controls, showed no deviation outside the normal range from the canonical medial-lateral organization of the ventral category-selective cortex (all |t(_23_)| values < 2.01; Figure 2J, see also spatial topography of category selectivity in each individual control in Figure S3).

#### Representational content per category

With the topography and spatial arrangement delineated, we then examined the extent to which the neural representations in each category-selective ROI in patients resemble those in TD controls and whether, within patients, this similarity differs for typically- versus atypically-sited ROIs (e.g., right VWFA in the three left resection patients). Using representational similarity analysis (RSA (65), see examples in Figure 3A-B and Methods for further details), we calculated, for each participant, the correlation between the preferred and non-preferred categories in each category-selective ROI (Figure 3C, purple vs. gray regions). Higher (Fisher-transformed) correlations reflect less differentiable representations and similar informational content, while lower correlations indicate more selective representations of the target category, respectively (Figure 3D).

**Figure 3.**
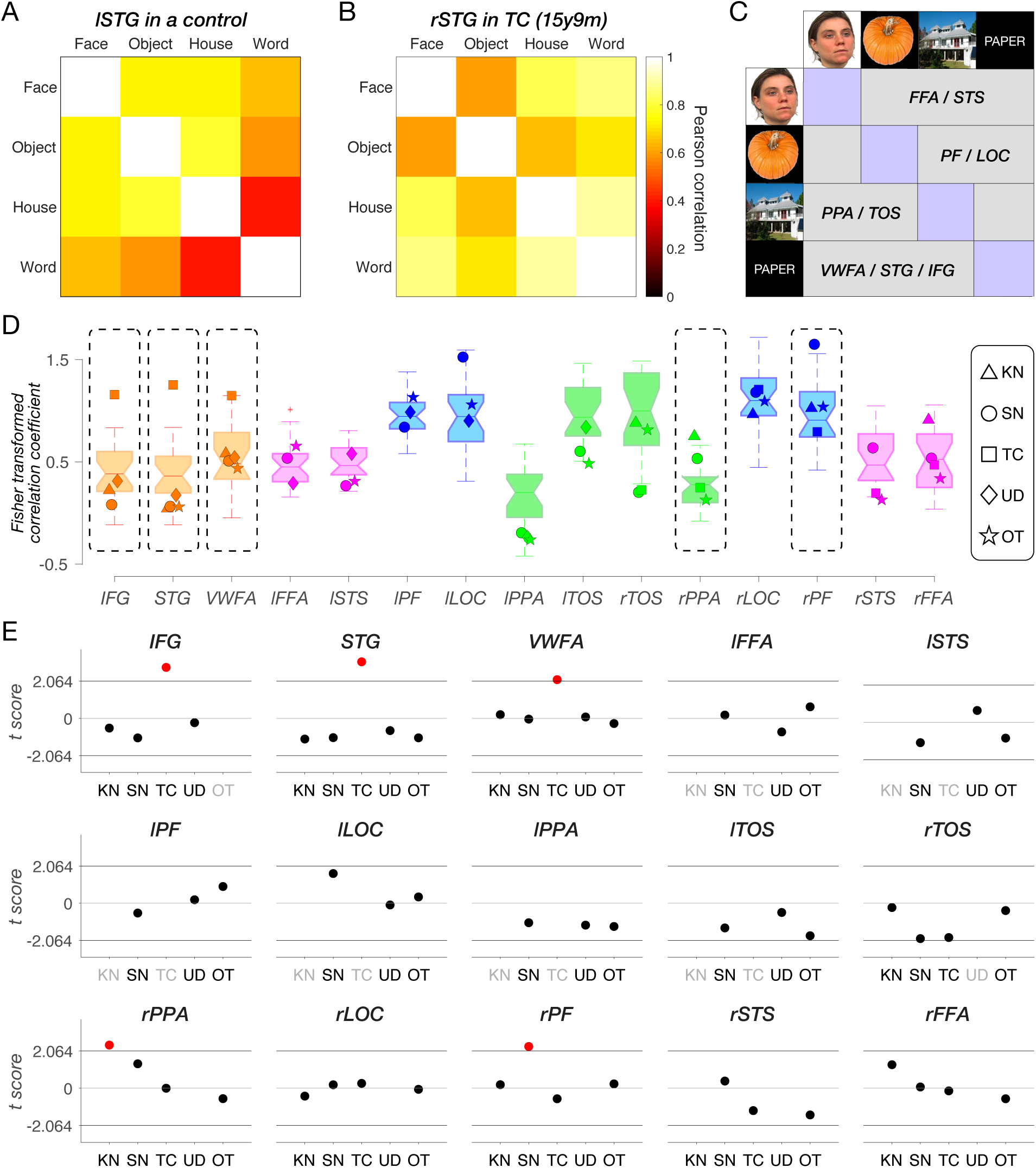
Representational similarity analysis of category-selective responses. STG = superior temporal gyrus, FFA = fusiform face area, STS = superior temporal sulcus, PF = posterior fusiform, LOC = lateral occipital complex, PPA = parahippocampal place area, TOS = transverse occipital sulcus, VWFA = visual word form area, STG = superior temporal gyrus, IFG = inferior frontal gyrus, l = left, r = right. **(A)** An example of left STG in a typically developing (TD) control, showing highly dissociable representation and low correlation between preferred (words) and non-preferred categories (faces, objects, and houses). **(B)** An example of right STG in TC (category localizer session 2), showing less dissociable representation and high correlation between the preferred (words) and non-preferred categories (faces, objects, and houses). **(C)** A schematic illustration of the representational similarity matrix in this analysis. For each ROI, the preferred category is depicted in purple, and all other categories are depicted in gray. **(D)** Fisher-transformed correlation coefficient between the preferred category and all other categories for each ROI in each patient’s last scan session and in TD controls. Each boxplot displays the full distribution of datapoints from the TD control group. A horizontal line inside the box indicates the median, the box represents the interquartile range between the first and the third quartiles, and the whiskers extend to the most extreme datapoints that are not considered outliers by the algorithm (MATLAB function: boxplot). Data points from each identifiable ROI in the patients are depicted with unique shapes per patient: triangle (KN), circle (SN), square (TC), diamond (UD) and star (OT). Details of the ROIs that are resected, not covered, or not found in the patients are shown in Figure S2. **(E)** Crawford’s t tests compared representational similarity in each identifiable ROI of patient scans to its respective TD control range. Red dots indicate significant deviations. Black x-axis labels indicate ROIs that can be defined either in the typical hemisphere or remapped to the opposite hemisphere. Gray x-axis labels denote ROIs that were resected, not covered, or not identified in the corresponding patients (see Figure S2).

We compared the correlation within each ROI in each patient against the corresponding correlations calculated within the same ROI in the TD control group using Crawford’s modified t test (Figure 3E). There were several cases of significantly less differentiable representation categories (higher correlations), relative to TD controls, in the LH resection patients: the atypical right IFG, STG, and VWFA in TC (orange squares in Figure 3D and red dots in Figure 3E, all remain significant following Benjamini-Yekutieli procedure (66) to control the false discovery rate (FDR) across multiple comparisons of 9 identified ROIs, at the adjusted first-, second-, and third-rank thresholds); the unilateral right PPA for KN (green triangle in Figure 3D and red dot in Figure 3E, no longer reached the adjusted first-rank significance threshold following Benjamini- Yekutieli procedure to control the FDR across multiple comparisons of 9 identified ROIs), and the right PF in SN (blue circle in Figure 3D and red dot in Figure 3E, no longer reached the adjusted first-rank significance threshold following Benjamini-Yekutieli procedure to control the FDR across multiple comparisons of 15 identified ROIs).

The same analysis applied to each individual control showed minimal deviation in representational structure with 6 out of 375 regions falling outside the normal range of the other 24 controls (Figure S4). These 6 deviating comparisons just marginally exceeded the threshold of the normal distribution, and, indeed, none of them survived the Benjamini-Yekutieli procedure to control the FDR across multiple comparisons of 15 identified ROIs in each control.

In other words, TC’s information content in the right VWFA and RH language areas (IFG and STG) differs from those regions in the LH of TD controls. In contrast, KN and SN, who also have RH-lateralized VWFA, IFG and STG regions due to LH resections encompassing the posterior VOTC, both have normal information content of these regions. Lastly, UD (the single RH resection patient) and OT (‘control’ patient, with LH resection outside VOTC) showed no differences in the representational structure compared to TD controls.

### Longitudinal analysis

Next, we present longitudinal data from three participants (TC, UD, and OT) who completed multiple neuroimaging sessions, following the format of the cross-sectional data but examining changes in metrics over time.

#### Spatial topography of category selectivity

As shown in Figure 4A-B, there were a small number of word-selective ROIs in TC and UD that emerge over time, i.e. they were not detectable on an earlier scan. This is especially evident in TC (left VOTC resection), in whom we first observed a right IFG (yellow) emerging at 13y11m, and a right STG (light orange) at 15y9m (Figure 4A). Neither region was detectable in TC’s first (pre- surgical) scan at 13y3m but her hospital records noted that language was lateralized to the LH. In TC’s separate postsurgical language localizer scan, the LH IFG were detected in a similar location using an established language localizer (Figure S8) (67).

**Figure 4.**
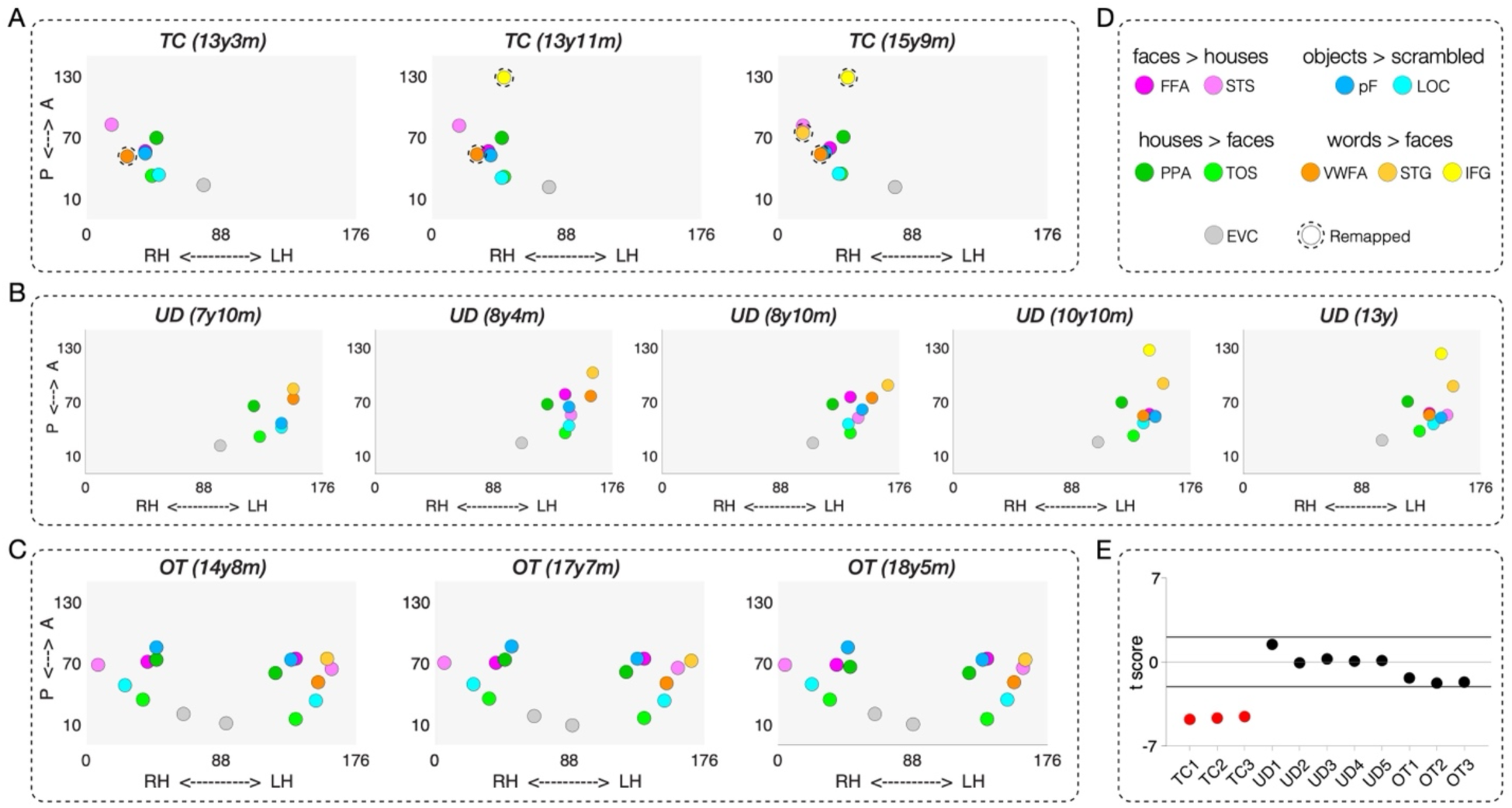
Spatial organization of category selectivity in each scan session in longitudinal patients TC, UD, and OT. (**A-C**) Category-selective topography across three scan sessions in TC, five scan sessions in UD, and three scan sessions in OT. The left side within each panel visualizes the average spatial distribution of category-selective regions of interest (ROIs). The x-axis represents coordinates in the medial-lateral direction for each hemisphere (LH: 88-176, RH: 0-88 in native space), and the y-axis represents coordinates in the anterior-posterior direction. Filled colored circles indicate ROIs that can be identified in a given scan, while circles surrounded by dotted lines represent ROIs for word function that are typically left lateralized but here are localized to the RH. **(D)** Contrasts to define category-selective activations for each region. **(E)** Crawford’s modified *t* score of difference in the spatial organization between each longitudinal patient and the TD control group.

For UD (right VOTC resection), the presence of the LH IFG was only visible in the last two sessions, as a result of a transition from partial to full brain coverage (Figure 4B). In UD’s presurgical clinical scan from the hospital, the LH IFG and LH STG were visualized and these very same regions were detected in our post-surgical language localizer scan (Figure S9) (67). For the pre- and post-surgical data and comparison, see Figure S1 of Liu et al. (46). All other ROIs were detectable across the five sessions in UD. Lastly, there were no changes in the number of identifiable ROIs across scans in OT (control anterior temporal resection; Figure 4C).

Next, we extended the medial-lateral analysis in Figure 2I-J to the longitudinal scans in TC, UD, and OT. Specifically, we observed a deviation in the spatial topography of ventral ROIs for all three of TC’s sessions (red dots in Figure 4E). By contrast, those for all five sessions for UD and three sessions for OT fall within the TD control distribution (black dots in Figure 4E). In other words, the reorganized RH VWFA in the case of posterior LH (but not anterior LH or RH) VOTC resection leads to a significant deviation in the spatial topography of category-selective ROIs that persists across time: both pre- and post-surgery for TC and longitudinally. Although TC’s remaining RH and UD’s remaining LH can accommodate both face and word representations, this differences between them in this analysis captures the more common bilateral face representation and left-lateralized word representation as the canonical category-selective topography in the TD controls, similar to those observed in the right-handed college-age students (32).

#### Representational content per category

Change in information content over time would manifest as a category that becomes more differentiable from other categories (increasing specificity) or that becomes less differentiable (diminishing specificity). As before, to estimate representational similarity, we calculated a Fisher- transformed correlation coefficient between the preferred category and all other categories for each ROI in each longitudinal scan session (Figure 5A). Patients’ correlation coefficients were then compared to the respective correlation coefficient distribution of the TD controls (Figure 5B).

**Figure 5.**
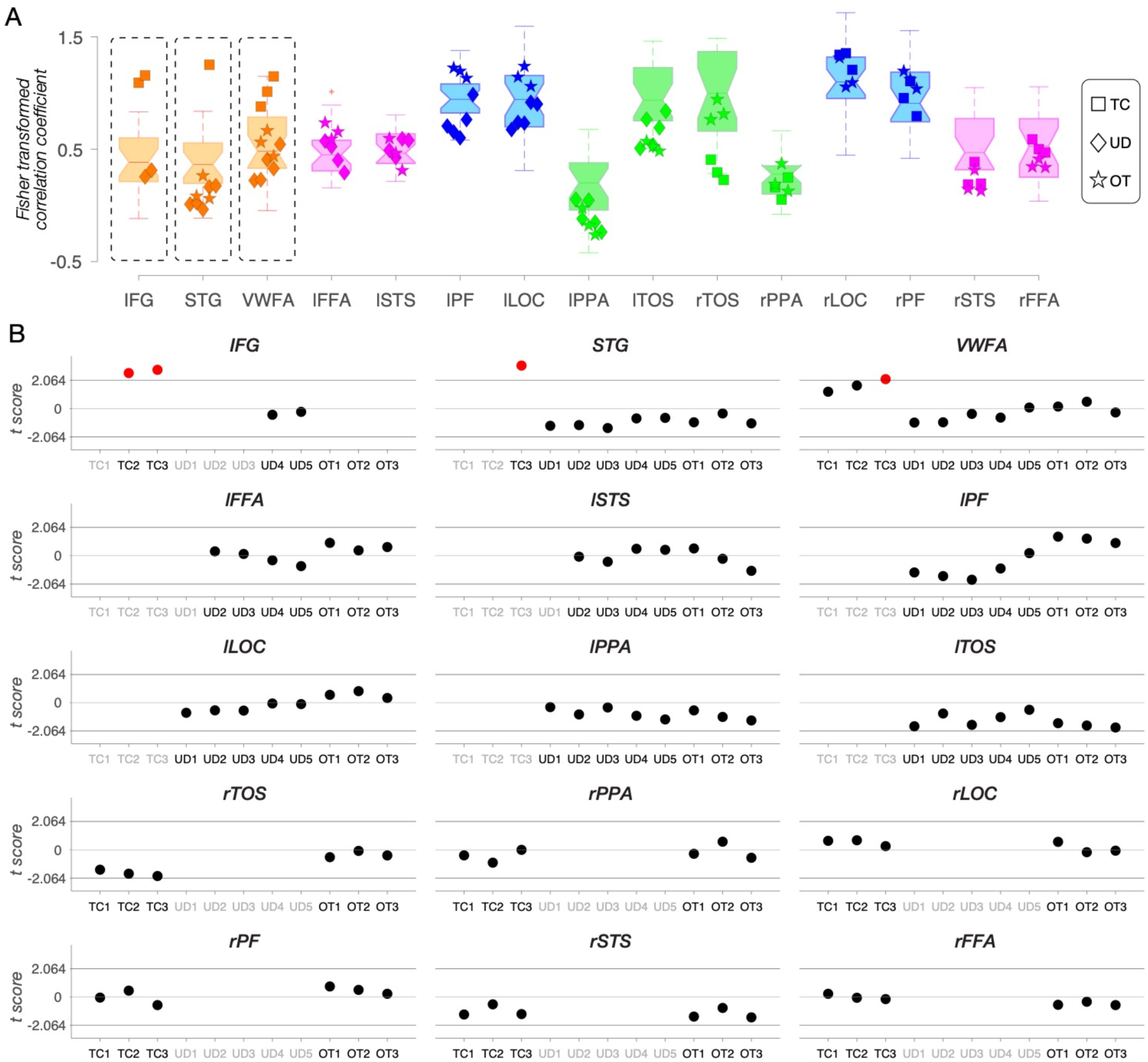
Representational similarity analysis of category-selective responses in each scan session in longitudinal patients TC, UD, and OT. **(A)** Fisher-transformed correlation coefficient between the preferred category and all other categories for each ROI in each longitudinal patient’s scan session and TD controls. Data points from each identifiable ROI in the longitudinal patients are depicted as squares (TC), diamonds (UD), and stars (OT). Details of the ROIs that are resected, not covered, or not found in the patients are in Figure S2. **(B)** Crawford’s *t* tests compared representational similarity in each identifiable ROI of longitudinal patient scans to the respective TD control range. Red dots indicate significant deviations. Black x-tick labels indicate ROIs that can be defined either in the typical hemisphere or remapped to the opposite hemisphere. Gray x-tick labels denote ROIs that were resected, not covered, or not identified in the corresponding patients (see Figure S2).

The longitudinal analysis of representational similarity confirms the stability of information content in UD (right VOTC resection) and OT (‘control patient’, anterior resection) over time (black dots in Figure 5B). In contrast, the information content for TC (left VOTC resection) differs from controls and across time (red dots in Figure 5B). Most notably, the information content of TC’s RH VWFA is within the TD range in the first 13y3m (pre-surgical) and second 13y11m (post-surgical) scan; however, by the third scan at 15y9m, the information content of right VWFA significantly deviates from TD controls (the red dot in Figure 5B; remained significant after applying the Benjamini-Yekutieli procedure (66) to control the FDR across multiple comparisons of 9 identified ROIs at adjusted third-rank threshold in CL3). Also, the right IFG (which only emerges on TC’s second and third scans) has information content outside the TD range (red dots in Figure 5B; remained significant after applying the Benjamini-Yekutieli procedure to control the FDR across multiple comparisons of 8 identified ROIs at the adjusted first-rank threshold in CL2 and 9 identified ROIs at the adjusted second-rank threshold in CL3). Similarly, the right STG (which only emerges on TC’s third scan) have information content outside the TD range (the red dot in Figure 5B; remained significant after applying the Benjamini-Yekutieli procedure to control the FDR across multiple comparisons of 9 identified ROIs at adjusted first-rank threshold in CL3).

#### Face and word representations in a single developing VOTC

In this final analysis, we zeroed in on regions of face- and word-selectivity and their relationship over time. Accommodating category-selective regions within a single posterior VOTC may be relatively straightforward for categories that typically have bilateral selectivity (e.g., PPA and LOC). The more pertinent question is how development with a single hemisphere comes to support categories, such as faces and words, that typically are lateralized by adulthood (albeit with faces generally less lateralized than words). More specifically, is there evidence of competition (68, 69) between face and word selectivity within the single preserved VOTC, and is this equivalent independent of which hemisphere is preserved?

To address this question, across multiple neuroimaging sessions within TC, UD, and OT, we scrutinized changes in face- and word-selective ROIs and contrasted these with changes in house- and object-selective ROIs. Specifically, we conducted both univariate and multivariate analyses with data drawn from an anatomically defined VOTC mask that encompassed the fusiform gyrus (FG) and the occipitotemporal sulcus (OTS), the anatomical regions for the categories of interest (cyan surface patches in Figure 6B, 6G, and 6L, also visible in volume space in Figures S5). For TC and UD, we examined the preserved VOTC. We also examined the LH FG/OTS over time in OT, the ’control’ patient with a left anterior resection. Because his word selectivity is strongly lateralized in the LH, similar to that in the TD controls, we chose to examine the LH, as more competition with face processing is expected there compared to the RH.

**Figure 6.**
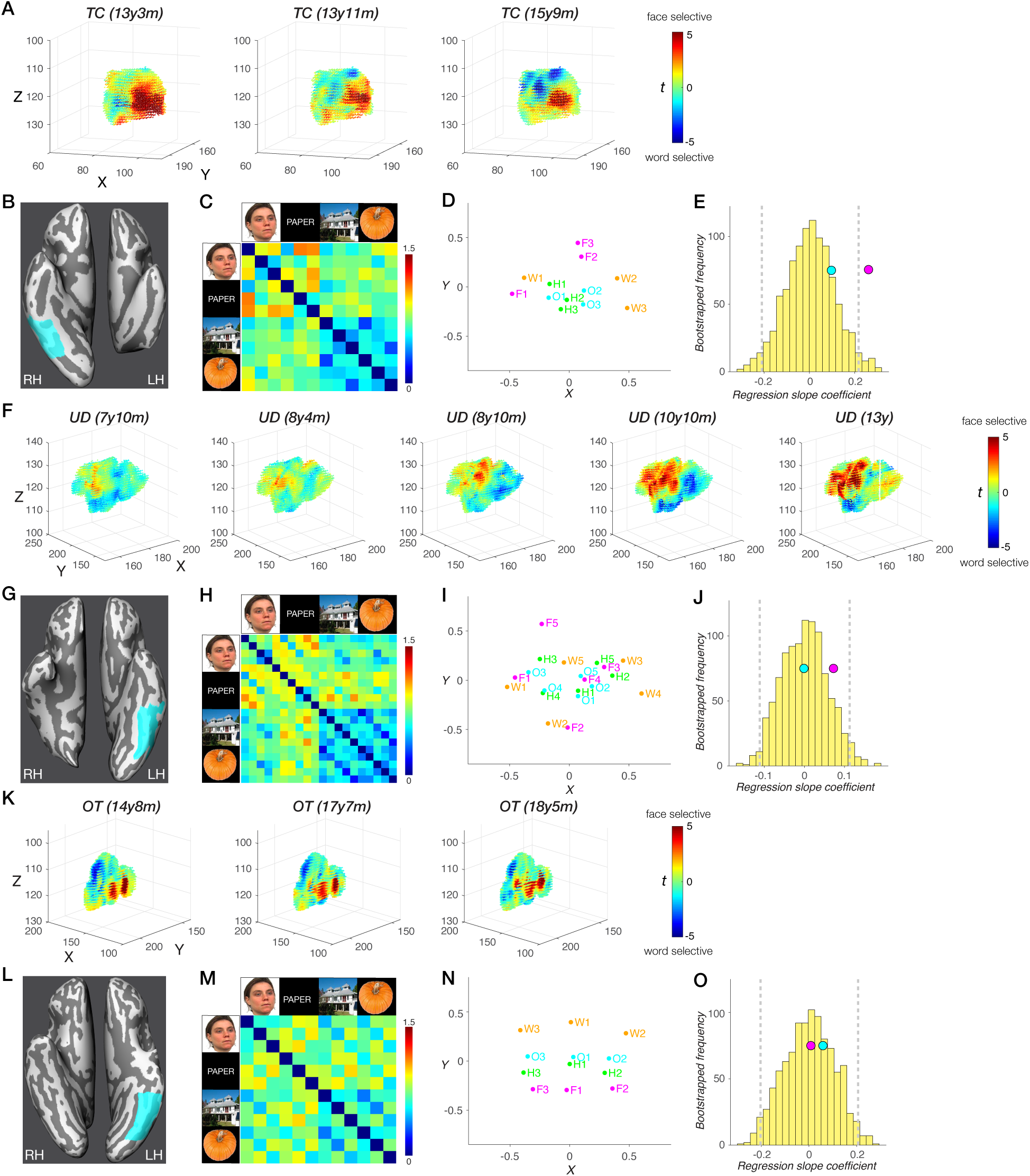
Changes in face and word representations over time in the anatomically defined fusiform gyrus/occipito-temporal sulcus (FG/OTS) in patients TC and UD, but not in OT. (**A, F, K**) Change over time in each voxel’s selectivity to faces over words within the FG/OTS region, where the XYZ coordinates (in native space) and *t*_(face-word)_ scores are plotted for each voxel. Higher selectivity to faces (dark red); higher selectivity to words (dark blue). We find significant differences in *t*_(face-word)_ scores between any two scan sessions in TC’s right FG/OTS and in UD’s left FG/OTS, except for the comparisons between scans 1 and 2. We find no significant differences in *t*_(face-word)_ scores between any two scan sessions in OT’s left FG/OTS. (**B, G, K**) FG/OTS (cyan) hand drawn in native surface space for TC (B), UD (G), and OT (K). see corresponding visualization in volume space in Figure S5. Total number of anatomical voxels (1mm isotropic) is 7307 in TC, 12428 in UD, and 12013 in OT. (**C, H, M**) Representational dissimilarity of category representations across sessions in TC’s right FG/OTS (C), UD’s left FG/OTS (H), and OT’s left FG/OTS (M). (**D, I, N**) Multidimensional scaling plot of category representations across sessions in TC’s right FG/OTS (D), UD’s left FG/OTS (I), and OT’s left FG/OTS (N). Words (orange), faces (magenta), houses (green), objects (blue). (**E, J, O**) A distribution of bootstrapped dissimilarity slopes (yellow histogram), face and word dissimilarity slope (pink circle), and house and object dissimilarity slope (cyan circle) as a function of the number of sessions in TC (E), UD (J), and OT (O). 95% CI (gray vertical dashed lines).

In TC, there was a significant increase in word-over-face selectivity over sessions across all 7307 voxels in right FG/OTS (more blue and fewer red voxels from scan 1 to 3 in Figure 6A; all |t| values > 6.522, all p values < 7.147e-11, two-tailed, independent samples t tests at the voxel level). This change was evident both in comparing TC’s pre-surgery to post-surgery scan (scans aged 13y3m to 13y11m) and thereafter across two post-surgery scans (scans aged 13y11m to 15y9m). In UD, a univariate analysis of the 12428 voxel of the left FG/OTS revealed clear increases in face-over-word selectivity over time (more red and fewer blue voxels from scan 1 to 5 in Figure 6F; all |t| values > 3.096, all p values < 0.002, except for the comparisons between scans 1 and 2, t(_24854_) = 0.197, p = 0.844, two-tailed, independent samples t tests at the voxel level). Finally, in OT, unlike in both UD and TC, there were no significant differences in face-versus word-selectivity between any two scan sessions (Figure 6K; all |t| values < 1.039, all p values > 0.299, independent samples t tests at the voxel level).

The quantifiable changes for faces and words in UD and TC contrast with the stable profile of object selectivity in FG/OTS over time. Specifically, in a univariate contrast between objects and scrambled objects, no significant increase or change between any two scan sessions were evident in TC (Figure S6, all |t| values < 1.549, all p values > 0.122) or in UD (Figure S7, all |t| values < 1.741, all p values > 0.082). The absence of change over time for objects in the context of changes in voxel selectivity for words and for faces in the single preserved VOTC for TC and UD indicates that not all categories are competing for representational space, thereby highlighting the specific competition between face and word representations. Taken together, our findings suggest that, longitudinally, there is competition between face and word representations for neural representational space within a single posterior VOTC (as observed in TC and UD).

However, this competition is not observed when bilateral posterior VOTC remains intact following unilateral anterior temporal lobe resection, as seen in OT.

These findings, which reveal changes in word and face selectivity in TC and UD, respectively, but not in OT, are highly suggestive of competition within the preserved VOTC. However, analysis of the distribution of t_(face-word)_ scores in the FG/OTS at each session does not indicate whether, over time, individual voxels within FG/OTS that were word-selective at one point in time become face-selective (or vice versa) at a later point in time, which would indicate competition for representation, as opposed to stable face- or word-selectivity within individual voxels across sessions.

To evaluate change in each voxel over time, we performed a McNemar’s test of change (with Yates’ correction) for each adjacent pair of sessions for each patient. Using the mean t_(face-word)_ scores across all sessions for each patient (OT, TC, and UD), we consistently applied a conservative criterion of t > mean + 1.5 for strong face selectivity and t < mean -1.5 for strong word selectivity to isolate those voxels with an initial strong commitment to a category (see Methods). We elected to focus on those voxels with strong selectivity as these should be least likely to change their category responsivity. If they did, however, this would be a clear demonstration of competitive dynamics and malleability. Within the 7307 voxels within TC’s right FG/OTS, there was a significant shift of voxel selectivity from strong face to word preference between each pair of adjacent sessions [CL1-2: McNemar X^2^ = 87.699, p < 0.001; CL2-3: McNemar X^2^ = 9.333, p = 0.002]. Likewise, amongst the 12428 voxels derived from UD’s mask, there were significant changes in strong face/word preference between each adjacent sessions in the first four sessions (McNemar X^2^ ranges from 8.10 to 21.061, all p values < 0.004) except for the last pair of sessions [McNemar X^2^ = 3.273, p = 0.070]. Interestingly, the saturation of responsivity to words versus faces in the last pair of sessions may reflect a stabilization of selectivity as UD reached age 13 years of age. Last, there were no significant changes in strong face/word preferences over time within a total of 12013 voxels in OT [CL1-2: McNemar X^2^ = 0.941, p values = 0.332; CL2-3: McNemar X^2^ = 0.563, p values = 0.453].

We next characterized changes in multivariate representations over sessions in TC, UD, and OT using representational dissimilarity matrix (RDM) (Figure 6C, 6H, 6M). The corresponding multi-dimensional scaling (MDS) plots are shown for TC in Figure 6D, UD in Figure 6I, and OT in Figure 6N. Each plot visualizes the similarity structure among stimuli as distances between conditions in a two-dimensional representation, which reveals more dispersed face (magenta) and word (orange) representations, compared to tighter clustering of object (blue) and house (green) representations. The greater separation between faces and words vs. houses and objects is consistent with a competitive dynamic in which representations diverge within the FG/TOS region. Next, we performed a bootstrapping linear regression analysis to derive an index of change between these pairs of representations over time. This was performed separately using the distance between faces and words, and between houses and objects, in each session in TC, UD, and OT.

In TC, the regression slope for face-word dissimilarity across three sessions (0.26, Figure 6E, pink circle) fell outside the 95% confidence interval (CI, [-0.210, 0.216]) of the bootstrapped null distribution (Figure 6E, yellow histogram), indicating increasing differentiation between face and word representations over development in the RH. In contrast, the regression slope for object- house dissimilarity across sessions (0.01, Figure 6E, cyan circle) fell within the 95% CI of the bootstrapped null distribution (Figure 6E, yellow histogram), suggesting a stable representation of houses and objects across sessions in right FG/TOS.

Across five sessions in UD, the regression slopes for faces and words (0.07, Figure 6J, pink circle) and for objects and houses (0.02, Figure 6J, cyan circle) fell within the 95% CI of the bootstrapped null distribution ([-0.102, 0.103]), indicating stable representations of all categories in his left FG/OTS region. However, we know from the voxel-wise McNemar tests of change that there may be some stabilization of category preference in UD’s last two sessions. If only the first 4 sessions are taken into account, UD’s face and word slope (0.12) is outside the bootstrapped distribution, consistent with the possibility of saturation of change in the voxel-wise analysis.

Last, across three sessions in OT, the regression slopes for faces and words (0.01, Figure 6O, pink circle) and for objects and houses (0.06, Figure 6J, cyan circle) both fell within the 95% CI of the bootstrapped null distribution ([-0.203, 0.206]), indicating stable representations for both in left FG/OTS.

Taken together, our findings suggest that, longitudinally, there are changes in extrastriate topography and representational content for the two patients with posterior VOTC resections (TC and UD) but not for the patient with anterior temporal lobe resection (OT). In the domain of face and word representations, the changes are clearest and are manifest as competition for neural representations. This competition is evident in TC following left VOTC resection and in UD right VOTC resection, as revealed in the voxel-wise analysis and from bootstrapping linear regression analyses that indexed the changes over time.

## Discussion

The goal of this investigation was to elucidate the nature of category-selective topography and representational similarity in human VOTC and the extent to which it is malleable. Given that the spatial organization of VOTC’s category-selective regions is highly replicable across individuals (20), one might predict rather minimal potential for change in VOTC aside from that associated with typical development. We have recruited individuals with unilateral childhood resection of VOTC (for the management of DRE), as all visual categories must presumably be accommodated within the preserved VOTC if these individuals are to maintain visual recognition behavior. As such, investigation of VOTC in such individuals will shed light on the potential for change in human ventral visual cortex. In the current work, we tracked changes in category selectivity and representational content in such individuals, using univariate and multivariate approaches, both cross-sectionally and longitudinally, to understand how a single VOTC comes to support various visual categories, some of which would ordinarily have been supported by the now-resected VOTC. We conducted further analyses on changes in areal selectivity for faces (FFA) and words (VWFA), as these have opposite stereotypical lateralization profiles, and the typical strong left-lateralization of words and its colocalization with language, in particular, poses a stringent test of plasticity when written words must be supported by the RH following left VOTC resection.

To address these issues, we acquired behavioral and neuroimaging data in three individuals with resections encompassing left VOTC (KN, SN, TC), one with right VOTC resection (UD), and one with a left anterior temporal lobe resection (OT) to serve as a ‘control patient’, with longitudinal imaging in TC, UD and OT. We also acquired data from 25 matched TD controls. Importantly, all patients performed within the range of TD controls on perceptual tests, except for KN (with the most extensive resection, a left hemispherectomy).

### Altered topographic profiles following cortical resection

In the patients, the patient control and TD controls, we identified, as far as possible, 17 ROIs, including category-selective regions as well as language areas and early visual cortex. The results indicated that spatial organization of category selectivity is flexible in patients with cortical resection versus controls, as evident, for example, by the emergence of lateralized language and word-selective regions in the typically non-dominant hemisphere (Figures 2 and 4). One noteworthy point is that, because the altered topographic profile was observed in those following smaller, lobar resection and not just following hemispherectomy suggests that it is the VOTC resection per se rather than the extent of the resection that determines the resulting topographic outcome. Also, noteworthy, is that a resection to the LH but situated more anteriorly in ventral cortex (in SN), leads to partial remapping with the preserved LH posterior VOTC still maintaining some signatures of typical topography, and an even more anterior temporal resection (in patient control OT) did not result in any topographical change of VOTC profile. These findings implicate the resection of the left posterior VOTC as the critical locus that triggers changes in topographic arrangement.

Some researchers have proposed that one strong constraint on VOTC topography concerns the medial-lateral arrangement of category-selective ROIs, specifically with the VWFA, with responses to written words activating a region that is more lateral than medial and that is proximal to laterally-situated regions that encode lexical and semantic information (63, 64, 70) — a spatial arrangement that is predictable even when measured prior to the acquisition of literacy (71, 72).

Our results suggest that this constraint may not be as strong as previously thought: only UD and OT had preserved medial-lateral topography, with the three left resection patients violating this constraint, presumably as a consequence of the atypical localization of the VWFA to the RH.

A further constraint on VOTC topography is thought to be the hemispheric lateralization of language. There is clear pressure, for the modal right handed individual, for the LH to develop orthographic representations proximal to, and thus co-lateralized with, language areas (24) so that the visual, phonological, and conceptual aspects of reading can be easily coordinated (2, 30, 73), and so that top-down language information can be integrated with bottom-up visual input (74–76). This pressure may explain why the asymmetry of word recognition in the left VWFA is greater than the more graded, bilateral profile of face recognition in the right FFA (32, 77).

Our results only partially uphold the colocalization assumption. The lateralization of word-selective cortex closely followed the lateralization of language in some but not all patients; all three patients with resections incorporating LH VOTC showed RH language regions (STG and IFG) and RH VWFA. In TC, however, as revealed in the longitudinal data, the RH VWFA was detected earlier than STG and IFG which could only be discerned in the second and third scans.

The RH co-localization of VWFA and language areas has also been reported previously in a case with LH resection (78), but there are also violations of this constraint, as illustrated by another case of RH lateralization of the VWFA but left-lateralization of language (79) and a further case of a VWFA in the absence of LH STS language region (80). Other atypical arrangements of the localization of the VWFA include abnormal recruitment of the anterior temporal lobes bilaterally for reading following left fusiform resection (81), the anterior shift of the VWFA within the LH (82), and even the presence of text-selectivity connected to LH motor and premotor regions via activity in left STS (83).

Beyond the topography, we were especially interested in understanding the representational content of category-selective regions, especially those in which topography deviated from the typical profile. As shown in Figure 3, the reorganization of word and language regions in TC to the right hemisphere was associated with less distinct representations of words in TC, but not in the RH VWFA of KN or SN, who also have LH resections, suggesting that VOTC category-selective areas within typical and atypical regions can maintain representations that are largely equivalent to those of the controls. It is also worth noting that word-related plasticity in TC appears to be underway pre-surgically (CL1, see Figures 4-5), although post-surgical changes are also detected and we return to this topic later in the Discussion.

### Dynamics of cortical plasticity

Our findings from the longitudinal data are particularly instructive in elucidating dynamic changes in the organization of higher-order visual cortex (Figure 4-6). We identified changes in spatial location of regions as well as in voxelwise selectivity across sessions in patients TC and UD, but not in the control patient, OT, whose longitudinal profile is remarkably stable (Figure 6). In both TC and UD, who have a left and right VOTC resection, respectively, changes in face or word selectivity were clearly evident across sessions (see Figure 6A and 6F), and the jockeying for representational space ultimately resulted in face- and word-selective voxels competing with and then abutting each other in a single VOTC. That both categories come to be situated in FG/OTS is consistent with the claim that these two categories require fine-grained foveal representations for the discrimination of their highly-similar exemplars and hence recruit the foveal-biased region of cortex (84). The changes and the asymmetry thereof for both word and face representations over time were confirmed by the multivariate analysis in which regression slopes for faces and words, but not objects and houses, across sessions fell outside of the bootstrapping null distribution for the preserved RH to a greater degree than the preserved LH.

### Implications for plasticity: which hemisphere and which areas accommodate new functions?

Some have argued that the functional and anatomical pressures that determine face and word selectivity arise from domain-specific innate constraints (85–87). The notion of a priori specifications of regional selectivity is difficult to reconcile with the flexibility and malleability of category- and content-specificity shown here. Given the opportunities for constructive remodeling or ‘recycling’ of VOTC, our findings also raise the question of exactly which cortical regions may be candidates for accommodating the VWFA or the FFA, if and when needed. Determining this is especially interesting for the VWFA in light of the relatively recent cultural adoption of word reading and the relatively late emergence of the VWFA ontogenetically (88–90).

Some have suggested that, during typical reading acquisition, face-selective regions can become word-selective (91, 92), and our findings here are consistent with this claim (as well as the reverse case in which face selectivity can be accommodated in word-selective cortex). Others have argued that regions that were limb-selective may be good candidates for visual word representations (21), and yet others have proposed that regions that are weakly selective and not committed to a particular stimulus category are possible sites too (18, 71, 93).

Our findings show that individual voxels that were initially strongly selective for one category—words or faces—can shift allegiance over time and become strongly selective for the other category. This evidence was more dramatic in TC whose VWFA had to be accommodated in the RH and abuts her FFA region, than in UD whose face-selectivity needed to be accommodated in his LH (which might have had a bias toward face selectivity in the first instance). Whether ‘recycling’ necessarily destroys another category-selective area in the course of its recycling or does so without destructive competition (93) is still debated. The findings here favor the latter: voxel allegiance shifts over time such that the representation of words adversely impact the representation of faces in the RH. Likewise, over time in the LH, voxels that are initially highly selective for word representations lose the competition and become more selective for face representations.

The pressure to reorganize the preserved hemisphere to accommodate face representations is likely weaker than for word representations, which are typically more unilateral. The FFA has precedence for more bilateral representation of function not only in adulthood, as noted above, but also early in development; for example, whereas before 24 months of age, either LH or RH damage can result in equivalent face recognition impairments (94), in adulthood, a lesion to the RH results in prosopagnosia more often and more severely than a LH lesion (95). In the context of language functions, which are also present in the preserved RH of our three left VOTC resection cases, bilateral underpinnings have also been reported, potentially consistent with claims of upregulation rather than major reorganization of cortex (96); in younger children, language appears to be activated bilaterally but, with age, the dominant LH appears to strengthen and just a ’weak shadow’ is detectable in the RH (97, 98) (for equivalent receptive vocabulary potential in the two hemispheres, see Liegeois et al. (99)).

This seemingly early bilateral pattern may account for the finding that, despite extensive resection, individuals with childhood hemispheric surgery averaged 85% correct for both word and face recognition, irrespective of whether the preserved hemisphere was the LH or RH (43, 44). Even in adulthood, a unilateral stroke to either hemisphere, however, results in a deficit in both face and word recognition, although to a greater degree for faces after RH stroke and for faces after LH stroke, suggesting some bilateral representation even in older individuals (100). Thus, following resection, amplification or up-regulation of a pre-existing function (96) may allow for the within-hemisphere enhancement of function rather than requiring interhemispheric transfer from a neurologically abnormal site.

### Pre- to post-surgical plasticity

Last, the current study examined whether the change of functional organization of VOTC was a result of the surgery or predated it. Most studies of DRE resection patients have only delineated VOTC categories post-surgery (37–40). One individual with a right occipital resection was shown not to evince any changes in pre- to post-surgical face selectivity following OTC resection, but this individual was 36-years-old (101). It is possible that, because of many years of presurgical seizure activity, changes may have occurred prior to surgery.

As part of our longitudinal investigation, in patient TC, there was no detectable selectivity for any category pre- or post-surgery in the ipsilesional LH, and all visual categories were uncovered in the contralesional RH (see Figure 2F), even for words which are typically represented in the LH in TD controls (Figure S3). Notably, word selectivity becomes increasingly prominent in the RH post-surgery (Figures 6A) and particularly so in language regions (Figure 5B): this emergence raises the possibility that surgery or seizure alleviation may have facilitated further plasticity of the contralesional hemisphere. Of relevance is that this expansion of word-selectivity was still ongoing in TC beyond age 13 years. It is also noteworthy that changes are detectable not only pre- to post-surgery, but that changes are appreciated across post-surgical scans; in other words, plasticity is not restricted to the direct effects of the surgery itself and longitudinal changes post-surgery still occur.

### Future directions

The findings of this combined cross-sectional and longitudinal investigation conducted with individuals with childhood resection for the management of epilepsy offer critical insights into the brain’s malleability during development. Focal epilepsy affects global brain-wide functional activity, beyond the site of the epileptogenic focus (102–104); as such, it is posited that, in cases of DRE or chronic epilepsy, persistent epileptic and interictal activity throughout development can result in progressively worse long-term negative cognitive outcomes (105–107). But this may not always be the case; the cortical visual system is apparently somewhat malleable and can be differently configured or upregulated for new functions. Indeed, despite the persistent homonymous hemianopia, the majority of post-surgical children have good visual outcomes (108). Additionally, epilepsy surgery appears to reverse the deleterious developmental effects of epileptic pathology (109). The cross-sectional findings here uncovered the categorical topography in VOTC, their spatial relationships and their information content, and revealed plasticity and spatial deviations, especially in the case of the VWFA (and associated language areas), although information content (representational structure) were chiefly similar to that observed in matched controls.

Also, although we tested visual function and competence and, with one exception, documented normal behavior, further investigation with even more fine-tuned behavioral assessment (69, 110) and more fine-tuned psychophysics might uncover other instances of deficient behavior. We have also limited our investigations to VOTC in those with childhood resections. Whether similar findings might emerge following other lobar resections and/or other cognitive functions remains to be investigated further. Answering these questions is important in furthering our understanding of cortical functional architecture and would also have direct translational and clinical implications.

## Materials and Methods

### Participants

Participants’ parents provided informed consent to participate in the protocol approved by the Institutional Review Boards of Carnegie Mellon University and the University of Pittsburgh (an interpreter assisted TC’s mother in completing the consent form), and participants provided assent. Participants were paid for their participation in the study.

#### Patients

Five right-handed pediatric patients who had undergone cortical resection (at University of Pittsburgh Medical Center Children’s Hospital of Pittsburgh) participated in this study. All were native English speakers except TC who came to and attended school in the United States from age 6. Table S1 lists the demographic and surgical information for each patient. Figure 1 includes the postoperative MRI as well as a detailed overview of the investigation (ages at behavioral testing and functional imaging using a category-selective localizer) for each patient.

KN and TC had a right homonymous hemianopia and UD had a left homonymous hemianopia, as determined by confrontation visual field testing and a 32-dot visual perimetry measure, with fixation enforced by eye tracking (111). SN and OT retained intact visual fields.

We were unable to obtain reliable pre- or post-surgical neuropsychological data from KN whose hemispherectomy was performed at 20 months or pre-surgical neuropsychological data from SN whose surgery was at 1 day of age but who is currently schooled in a regular age- appropriate classroom setting. Intelligence quotient scores for TC could not be obtained presurgically as her English skills were not sufficiently well-developed at that time (although she was in a regular high school at the time of this testing). UD’s presurgical IQ scores were at least 1 SD above the standard mean of 100, and little change was evident from pre- to post-surgery. OT had a presurgical IQ of 122 and a postsurgical IQ of 127, with academic skills and performance above age and grade expectations. See Table S2 for additional information obtained from neuropsychological investigations for each patient.

#### Controls

Twenty-five age-matched TD controls (all right-handed, ten females, average age at scan: 12±3 years, see Table S3 for their ages at scan), with normal or corrected-to-normal vision and no neurological history, participated in the fMRI studies. Four of the controls (right-handed, 2 males) also participated in the behavioral testing session, and we recruited an additional 17 controls (right-handed, two males) for behavioral testing to obtain a distribution against which to compare the visual perception performance of the patients. Out of the 21 behavioral controls, one did not participate in the object-matching task.

### Behavioral experiments

In all patients, intermediate-level vision (contour integration and Glass pattern) and high-level pattern recognition (face and object recognition) were assessed using a 14" Dell laptop with viewing distance of roughly 60 cm. The contour integration, Glass pattern, and object matching tasks in controls were performed using the same laptop as in patients.

#### Contour Integration

The contour integration task used two collinearity conditions (target Gabor elements had either ± 20° or ± 0° collinearity) (49). Participants were instructed to use the keyboard to indicate whether an embedded egg-like shape pointed to the left or right (Figure S1A). Background Gabor elements were varied according to a one-up (after a wrong response), three-down (after three correct responses) staircase procedure, and the experiment continued until ten reversals in the staircase occurred. The threshold score reported in Table 1 was calculated from the geometrical mean spacing of the final 6 reversals. The overall area covered by all the Gabor elements extended about 17.6° horizontally and 12.6° vertically.

#### Glass Patterns

The perception of shape or global form was assessed using thresholds derived from a glass pattern (112). In this task, we varied the percentage of signal dots using a one-up (after an incorrect response), three-down (after three correct responses) adaptive staircase method to measure the 75% threshold for detecting the concentric swirl (50) (Figure S1B). The staircase started at 95% signal and terminated after 10 reversals. The threshold was measured from the geometric mean of the last 6 reversals.

#### Face recognition

We used the Cambridge Face Memory Test for Children (51) and followed the standard test instructions (see Figure S1C). Participants studied 5 faces and then, in subsequent trials, identified each ’old’ face from amongst new, distractor faces. The test was conducted using upright and inverted faces, in separate blocks. There were 60 trials in each orientation consisting of 15 introductory trials, 25 trials without noise, and 20 trials with added noise. Performance was the percent correct out of all 60 trials, separately for upright and inverted faces. The patients’ performance was compared to the control group from Croydon et al. (51), 10-year-olds, N = 41.

#### Object recognition

All controls and patients, except for KN, underwent testing for object recognition using an object judgment task adapted from (52). In this task, two objects were presented simultaneously—one above and one below the midline largely to circumvent the hemianopia—for same/different discrimination. The task consisted of 100 trials, 40 same and 60 different (twenty per difference level), randomly intermixed. When the objects differed, they could differ at the basic (e.g., duck vs. vehicle), subordinate (e.g., chair vs. piano), or exemplar level (e.g., table1 vs. table2), reflecting increasing perceptual similarity. The display remained on the screen until the participant’s response, with one key indicating ’same’ and another ’different’." Instructions encouraged both speed and accuracy (and both were measured), and a 25-trial practice block familiarized the participant with the task.

Patent KN was tested on the Cambridge Bicycle Memory Test for Children (53).

Participants are instructed to study a set of bicycles and then identify these amongst novel images of bicycles. Following standard instructions, 72 trials are presented (learning stage: 18 trials; test stage with novel viewpoints: 30 trials; test phase with noise overlaid: 24 trials). The scores were converted to percent correct out of all 72 trials, with separate calculations for upright and inverted bikes. The performance of the age-matched control group was determined using the data from Bennetts et al. (53), UK school year=6 (age 11), N = 22.

### fMRI experiments

#### MRI setup

MRI data were acquired on either a Siemens Verio 3T magnet at the Scientific Imaging and Brain Research Center or a PRISMA at the Carnegie Mellon University-Pitt Brain Imaging Data Generation & Education Center (RRID:SCR_023356), using a 32-channel phased array head coil. The patients had been scanned previously at the UPMC Children’s Hospital of Pittsburgh as part of their clinical examination and were comfortable in the magnet.

#### Structural MRI

A high-resolution (1mm^3^ isotropic voxels, 176 slices, acquisition matrix = 256 × 256, TR = 2300 ms, TE = 1.97 ms, inversion time = 900 ms, flip angle =°9, acceleration/GRAPPA = 2, scan time = 5min 21s) T1-weighted whole brain image was acquired for each participant using a magnetization prepared rapid gradient echo (MPRAGE) imaging sequence for localization, co- registration, and surface reconstruction purposes.

#### Functional MRI

In patient UD and OT, and for two TD controls, fMRI data were collected with a blood oxygenation level-dependent (BOLD) contrast sensitive echo planar imaging (EPI) sequence (TR = 2000ms, TE = 30ms, voxel size = 2.5mm^3^, interslice time = 79ms, flip angle = 79^°^, acceleration/GRAPPA = 2, 27 slices). In the other three patients (KN, SN and TC) and 23 matched controls, fMRI data with whole brain coverage (69 slices) were collected with a multiband acceleration factor of 3 and voxel size = 2 mm^3^ (all else equal to standard protocol). For all participants, slice prescriptions were AC-PC aligned.

#### fMRI task and stimuli

The visual presentations were generated using MATLAB (The MathWorks, Natick, MA) and Psychtoolbox (www.psychtoolbox.org). Images were back-projected onto a screen in the bore of the scanner. A trigger pulse from the scanner synchronized the onset of the stimulus presentation to the beginning of the image acquisition. During the category localizer tasks, a central fixation dot remained on the screen to orient participants’ fixation (see Figure 2A). Participants were instructed to maintain fixation, and eye movement was monitored to enforce fixation using an ASL eye tracker (Applied Science Laboratories, Billerica, MA) or an EyeLink 1000 (SR Research, Ottawa, Canada).

In each session, participants completed three runs of the fMRI category localizer task (45, 46, 113). The functional runs adopted a block design with stimuli from five categories (Figure 2A): faces (from the Face Place dataset (114)), houses, objects, scrambled objects, and words.

Each run consisted of 3 repeats of each category (8 TRs, 16 images) in pseudorandom order with a fixation baseline (4 TRs) between all conditions. Thus, each run contained 15 categories and 16 fixation baselines and lasted 6min 8s (184 TRs). Participants detected an immediately repeating image (one-back task) via an MR-compatible button glove using their index finger, and there was a single repeat per block. This response instruction was designed to engage participants maximally while keeping the task relatively easy for the children (overall accuracy: 95.8±3.2%). In the two longitudinal VOTC cases, TC and UD, a post-surgical (functional) language localizer was acquired (67). We used a block design with two categories: sentences and nonword strings. Participants were instructed to press one button (index finger) to indicate if the blue word/nonword shown immediately after the sequence (9 words/nonwords) matched one of the words/nonwords in this sequence, and another button (middle finger) to indicate a non-match.

This response instruction was designed to maximally engage participants while keeping the task relatively easy. Standard general linear model (GLM) analyses were run with 3 predictors (sentences, nonword strings, fixations), each convolved with a canonical hemodynamic response function (115). Language-selective ROIs were determined using the sentences-nonwords or sentences-fixation contrast. Using this task, we confirmed the left hemisphere (left IFG) dominance in both TC and left STG activation in TC.

### fMRI Data Analysis

#### Preprocessing

Preprocessing of the anatomical MRI included brain extraction/skull stripping, intensity inhomogeneity correction, and AC-PC alignment. Given the variability in the extent and site of the lesions in the patients, there was no spatial normalization, and analyses were conducted in native space. Functional data were 3D-motion corrected (trilinear/sinc interpolation), slice-time corrected, and temporally filtered (high-pass GLM Fourier = 2 cycles). Functional runs were co-registered with the structural scan using boundary-based registration approach. To permit the multivariate analysis, no spatial smoothing was applied.

To ensure accurate within-subject comparison in the longitudinal patients, we co-registered all functional runs in each patient to the structural MRI from the first category localizer session and carefully monitored the head motion and the temporal signal-to-noise ratio (tSNR) across sessions (see tSNR equation). Despiking of high-motion time points in TC and UD was performed using the ArtRepair toolbox (116) in Statistical Parametric Mapping (https://www.fil.ion.ucl.ac.uk/spm/).

#### Head motion

During each run, for each participant and control, the head motion was calculated from the combination of three translation parameters (in millimeters) and three rotation parameters (in degrees) using the following equations:

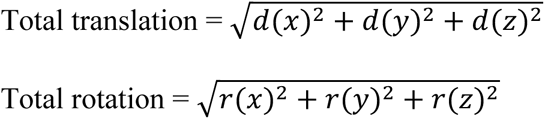

The average head motion for patients and controls was very similar: for patients, it was 0.44±0.20 mm (translation) and 0.48±0.28 degrees (rotation), and for controls it was 0.43±0.23 mm (translation) and 0.47±0.26 degrees (rotation).

#### Temporal signal-to-noise ratio

To ensure comparable fMRI data quality across participants as well as within-participant across sessions, we used tSNR as an index of the temporal SNR for each voxel. To minimize the influence from signal dropout due to resection, we excluded those voxels in the lesioned brain region (Figure 1, left) from the tSNR calculation in each patient. For each run, tSNR was calculated as the mean signal of the fMRI time series divided by the standard deviation of the noise in the time series: SNR_(temporal)_ = *μ* _time series_ ***/***σ _time series._

#### General linear model

For each run, a standard general linear model was performed. The regressor for each condition (faces, houses, objects, scrambled objects, and words) was defined as a boxcar function convolved with a canonical hemodynamic response function (*115*). To avoid overfitting, fixation conditions were not included.

#### Region of interest (ROI) definition

A total of 17 ROIs were identified using a set of contrasts. In each participant, category-selective ROIs were defined as a sphere (radius: 7mm) centered on the peak voxel under each paired contrast (see below, same as the method used in Liu et al. (45)).

The FFA (54, 55) was defined as the region in the mid-fusiform gyrus with greater activation for faces compared with houses (magenta in Figure 2). The STS (56) was defined as the region in the posterior STS with greater activation for faces compared with houses (pink in Figure 2). The pF (60, 61) was defined as the posterior bank of the fusiform gyrus with greater activation for intact objects compared with scrambled objects (dark blue in Figure 2). The LOC was defined as the region on the lateral bank of the fusiform gyrus extending dorsally into the middle occipital gyrus (below the lateral occipital sulcus) with greater activation for intact objects compared with scrambled objects (light blue in Figure 2). The PPA (57) was defined as the region in the anterior portion of the parahippocampal gyrus with greater activation for houses compared with faces (dark green in Figure 2). The transverse occipital sulcus (TOS) (58) was defined as the region in the TOS with greater activation for houses compared with faces (light green in Figure 2). The VWFA (62) was defined as a region in the left or right VOTC with greater activation for words than faces (dark orange in Figure 2). The STG (commonly known as Wernicke’s area) was defined as a region in the left or right posterior part of the STG with greater activation for words than faces (light orange in Figure 2). Last, the IFG (commonly known as Broca’s area) was defined as a region in the left or right inferior frontal gyrus (yellow in Figure 2).

#### The spatial relationship between ROIs

As a means of estimating the extent to which the spatial organization of the different ROIs was preserved in the patients and the possibility of change over the multiple within-subject sessions, we first extracted the native x and y coordinates of the peak voxel in each identifiable ROI for each participant (Figure 2C-H and Figures S3-4). We elected to stay in the native space for this analysis because we were unable to normalize the lesioned brains without further distortion.

Next, we quantified potential deviations of the medial-lateral organization principle of the ventral visual pathway by correlating (using MATLAB function corr) the x coordinates of all identifiable ventral ROIs (from medial to lateral: EVC—PPA—pF—FFA—VWFA) in each patient with the average x coordinates of these ROIs obtained for the controls. We then used Crawford t test to evaluate whether a patient’s coordinates fell outside of the normal distribution (Figure 2I). We also applied Crawford t test in each control to evaluate whether a control’s coordinates fell outside of the distribution of rest of the controls.

#### Representational structure of category selectivity

We applied RSA (65) to characterize the nature of the representations within each ROI. We computed Pearson correlation coefficients across all categories (face, object, house, and word) based on the beta value for all voxels in each ROI (see examples in Figure 3A-B). We then applied Fisher transformations to permit the use of parametric statistics. Finally, for each ROI, we calculated the average correlation between the preferred category (Figure 3C, purple regions) and all other categories (Figure 3C, gray regions). With FFA/STS as an example, the preferred category is faces, and the non-preferred categories include objects, houses, and words (Figure 3C). High (Fisher transformed) correlation coefficients reflect less selective representations, whereas low (Fisher transformed) correlation coefficients reflect more dissociable or unique representations of the preferred category (Figure 3D).

#### Multi-dimensional scaling

A multi-dimensional scaling (MDS) algorithm was run on the dissimilarity values stored in the upper (or equivalently the lower) triangle of the RDM. The resulting MDS plot visualizes the similarity structure coded in the RDM as distances between conditions in a two-dimensional representation (Figures 6D, 6I, and 6N).

### Statistical Analysis

#### Crawford’s modified t-test

We adopted a matched case-control design to compare the findings from each individual patient to their matched controls using modified t-tests (117) for both behavioral and fMRI experiments. The α criterion for all tests was .05, with Benjamini-Yekutieli procedure applied to control the false discovery rate (FDR) across multiple comparisons (66).

#### McNemar test of change

We applied McNemar’s test of change (with Yates’ correction) to evaluate change from face to word selectivity and word to face selectivity in each voxel in FG/OTS in TC, UD, and OT between each pair of adjacent sessions. Using the mean *t*_(face-word)_ scores from each session for each patient (OT, TC, and UD), we consistently applied a conservative criterion of t > mean + 1.5 for strong face selectivity and t < mean -1.5 for strong word selectivity to isolate those voxels with an initial strong commitment to a category.

Out of a total of 7307 voxels within TC’s right FG/OTS, 2424 voxels are strongly face- selective and 1205 voxels are strongly word-selective in CL1, 700 voxels are strongly face- selective and 1354 voxels are strongly word-selective in CL2, and 1097 voxels are strongly face- selective and 2157 strongly word-selective in CL3. From CL1 to CL2, 121 voxels shifted from face- to word-selective, and 12 shifted from word- to face-selective. Between CL2 and CL3, 18 voxels changed from face- to word-selective, and 3 from word- to face-selective.

Out of the 12428 voxels derived from UD’s FG/OTS, 834 voxels are strongly face- selective and 2610 voxels are strongly word-selective in CL1, 438 voxels are strongly face- selective and 1510 voxels are strongly word-selective in CL2, 1336 voxels are strongly face- selective and 2230 voxels are strongly word-selective in CL3, 3781 voxels are strongly face- selective and 1483 voxels are strongly word-selective in CL4, and 3292 voxels are strongly face- selective and 1840 strongly word-selective in CL5. From CL1 to CL2, 82 voxels shifted from face- to word-selective, and 32 shifted from word- to face-selective. Between CL2 and CL3, 0 voxels changed from face- to word-selective, and 10 from word- to face-selective. Between CL3 and CL4, 1 voxel changed from face- to word-selective, and 15 from word- to face-selective.

Between CL4 and CL5, 9 voxels changed from face- to word-selective, and 2 from word- to face- selective.

Out of the 12013 voxels in OT’s FG/OTS, 2082 voxels are strongly face-selective and 2372 voxels are strongly word-selective in CL1, 1471 voxels are strongly face-selective and 1685 voxels are strongly word-selective in CL2, and 1255 voxels are strongly face-selective and 1313 strongly word-selective in CL3. From CL1 to CL2, 11 voxels shifted from face- to word- selective, and 6 shifted from word- to face-selective. Between CL2 and CL3, 6 voxels changed from face- to word-selective, and 10 from word- to face-selective.

#### Bootstrapping linear regression

We derived a regression slope as an index of change to capture the relationship between face and word, or house and object representation over time. In TC and OT, bootstrapped regression slopes were calculated from the randomly picked 3 values (as a proxy for 3 sessions in TC/OT) after shuffling the condition labels in the upper (or equivalently the lower) RDM 1000 times in Figure 6C (TC) or Figure 6M (OT). This procedure yielded a distribution of the bootstrapped regression slopes (yellow histogram in Figure 6E and 6O), and the face and word dissimilarity slope (pink circle in Figures 6E and 6O) and the house and object dissimilarity slope (cyan circle in Figures 6E and 6O) was each compared with the 95% CI of the bootstrapped null distribution (Figure 6E and 6O, gray vertical dashed lines).

We performed a bootstrapping linear regression analyses in UD in which the condition labels in the upper (or equivalently the lower) RDM in Figure 6H 1000 times were shuffled and 5 values (as a proxy for a total of 5 sessions in UD) randomly picked each time to obtain the bootstrapped regression slope distribution (Figure 6J, yellow histogram). To establish the statistical significance of the difference between bootstrapped slopes and the face and word dissimilarity slope (pink circle) or the house and object dissimilarity slope (cyan circle), we calculated the 95% CI of the bootstrapped null distribution (Figure 6J, gray vertical dashed lines). We note that we have previously reported data for the first 4 sessions in UD (46) but have extended the data set here and recalculated the distribution.

## Acknowledgments

We thank the participants and their families for their time and cooperation; Scott Kurdilla, Mark Vignone, and Debbie Viszlay for their help in acquiring the imaging data; and Drs. Nicholas Blauch, Carl Olson, Michael Tarr, and the VisCog group at Carnegie Mellon University for the fruitful discussions. Face images for category localizer courtesy of Michael J. Tarr, Carnegie Mellon University, http://www.tarrlab.org/; funding provided by NSF award 0339122.

## Author contributions

TTL: Conceptualisation, Methodology, Software, Validation, Formal analysis, Investigation, Data Curation, Writing - Review & Editing, Visualisation, Project administration

MCG: Conceptualisation, Methodology, Software, Formal analysis, Investigation, Data Curation, Writing - Review & Editing, Project administration

AMSM: Methodology, Software, Investigation, Editing SR: Methodology, Software, Investigation, Editing JZF: Methodology, Software, Editing

CP: Patient recruitment and management, Investigation, Editing DCP: Conceptualisation, Writing, Editing

MB: Conceptualisation, Methodology, Funding acquisition, Supervision, Data interpretation, Writing, Editing.

## Data and materials availability

### Data availability

The dataset will be freely and publicly available upon publication on the Carnegie Mellon University data repository KiltHub (Figshare) at doi: 10.1184/R1/24898245. All data are available in the main text or the supplementary materials.

### Code availability

E-prime (Psychology Software Tools, Inc., PA), MATLAB 2016b (MathWorks, MA), and Psychtoolbox (www.psychtoolbox.org) were used to present the stimuli. A combination of publicly available software packages (Freesurfer, SPM) and commercial software (BrainVoyager, Matlab) and were used for fMRI preprocessing and analysis. Customized code, source behavioral and fMRI data, and high-resolution figures are available on Github (https://github.com/tinaliutong/VOTC-plasticity).

