## Supplementary Materials for "Cross-sectional and longitudinal changes in category-selectivity in visual cortex following pediatric cortical resection"

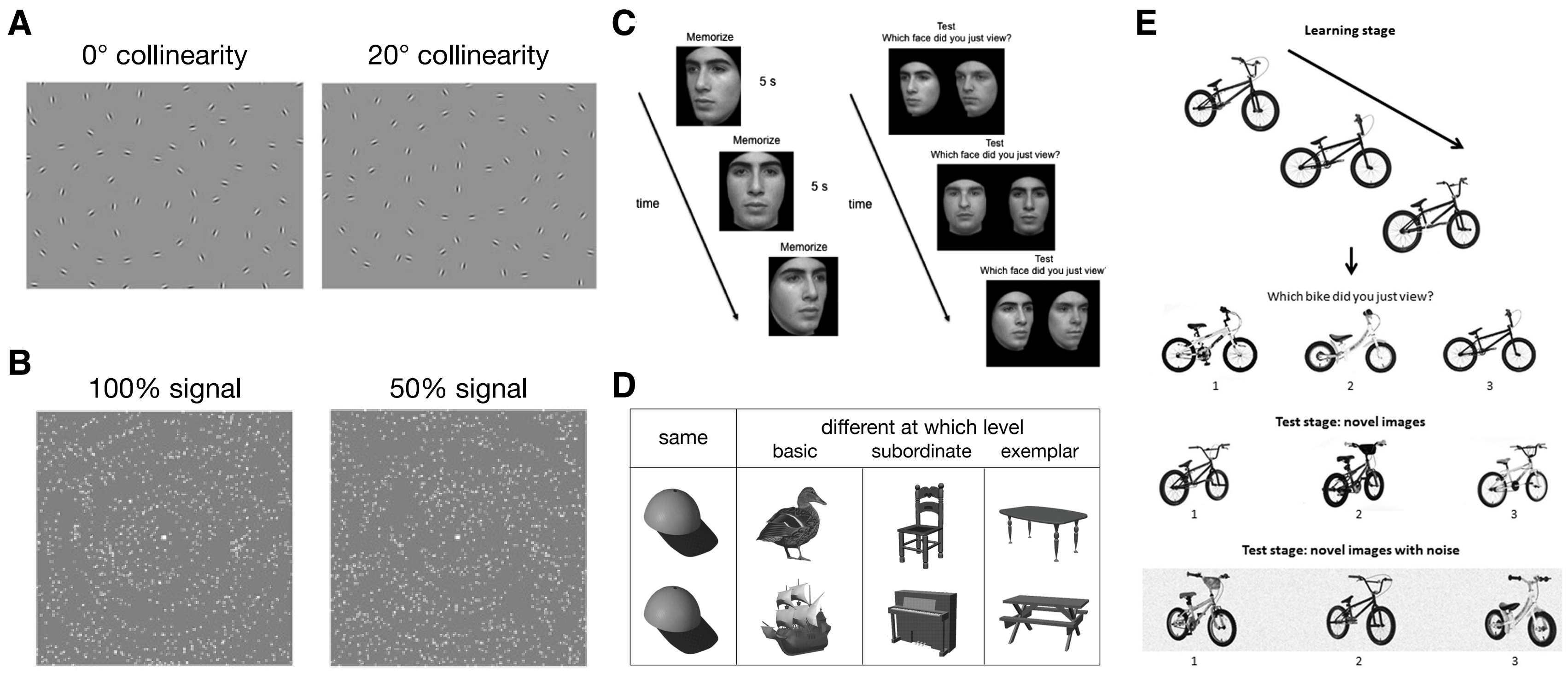


**Fig. S1. Design of behavioral experiments.**
(**A**) Contour integration (Hadad et al., 2010). Task: Participants viewed a brief presentation of either aligned or misaligned Gabor patches (in separate blocks) and indicated the leftward or rightward direction of the embedded 'egg-like' shape. (**B**) Glass pattern (Lewis et al., 2002). Task: Participants indicated which of the two displays had a more concentric swirl. (**C**) Cambridge Face Memory Test for Children (Croydon et al., 2014). Task: Participants were instructed to remember target faces and subsequently identify them amongst an array of distractor faces. (**D**) Object matching experiment (Gauthier et al., 1999). Task: Participants made same/different discriminations on pairs of objects and pressed a 'same' or 'different' button to indicate their response. (**E**) Cambridge Bicycle Memory Test for Children (Bennetts et al., 2017). Task: Participants were instructed to remember target bicycles and subsequently identify them amongst novel images of bicycles.

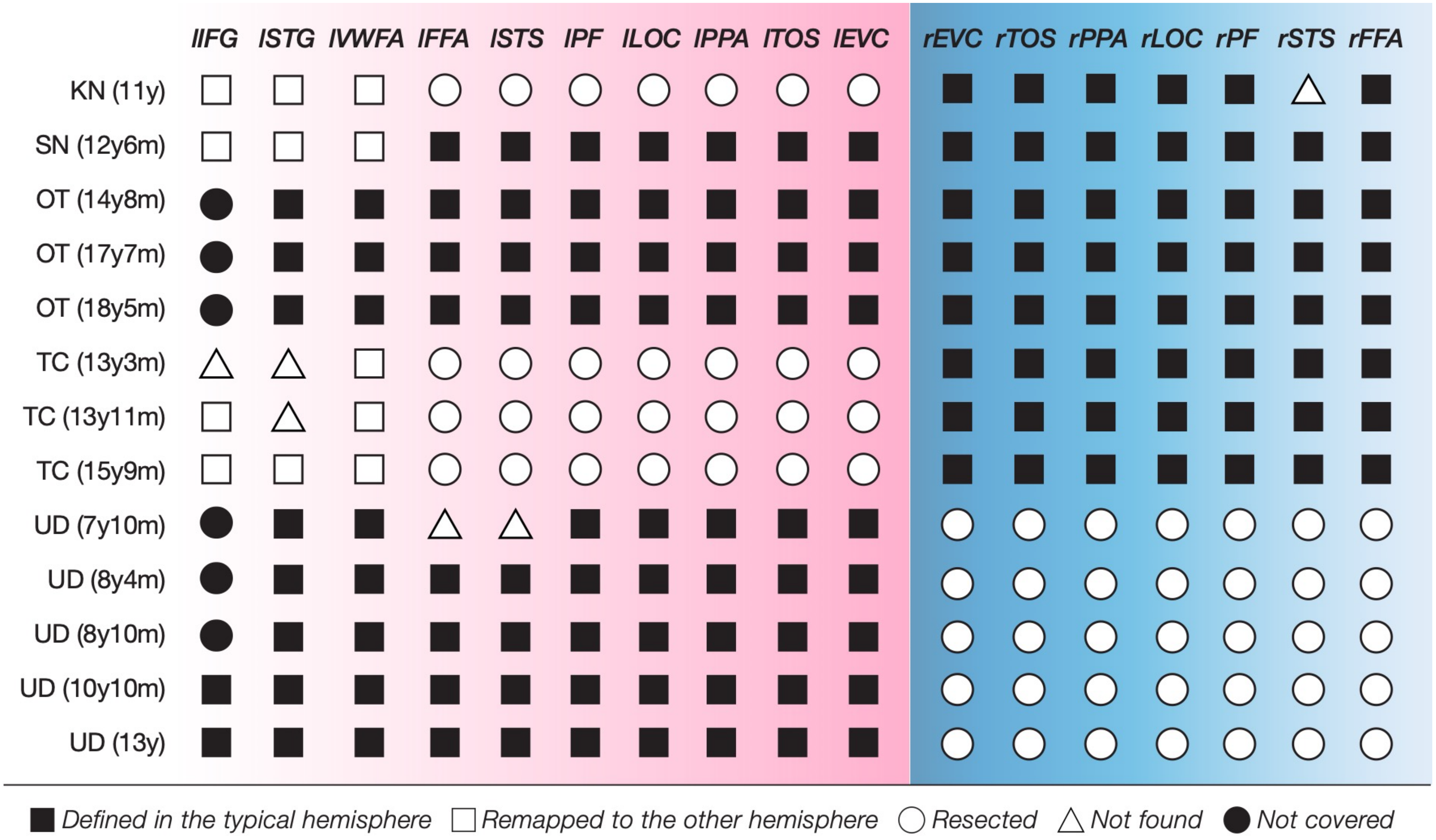


**Fig. S2. The presence of category-selective regions of interest (ROIs) in each patient.**
l = left, r = right, IFG = inferior frontal gyrus, STG = superior temporal gyrus, VWFA = visual word form area, FFA = fusiform face area, STS = superior temporal sulcus, PF = posterior fusiform, LOC = lateral occipital complex, PPA = parahippocampal place area, TOS = transverse occipital sulcus, EVC = early visual cortex.

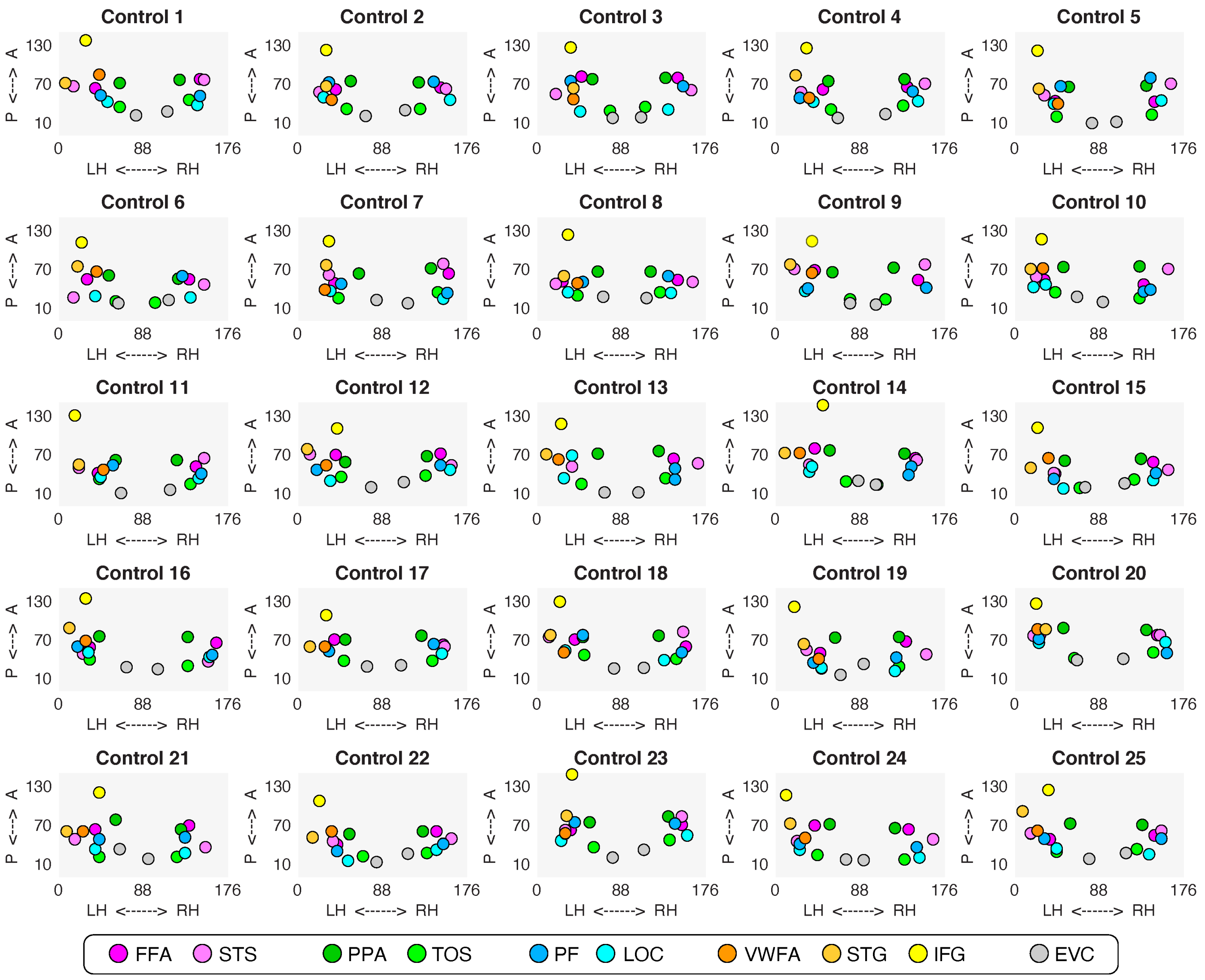


**Fig. S3. Spatial organization of category selectivity in each individual control (n=25).**

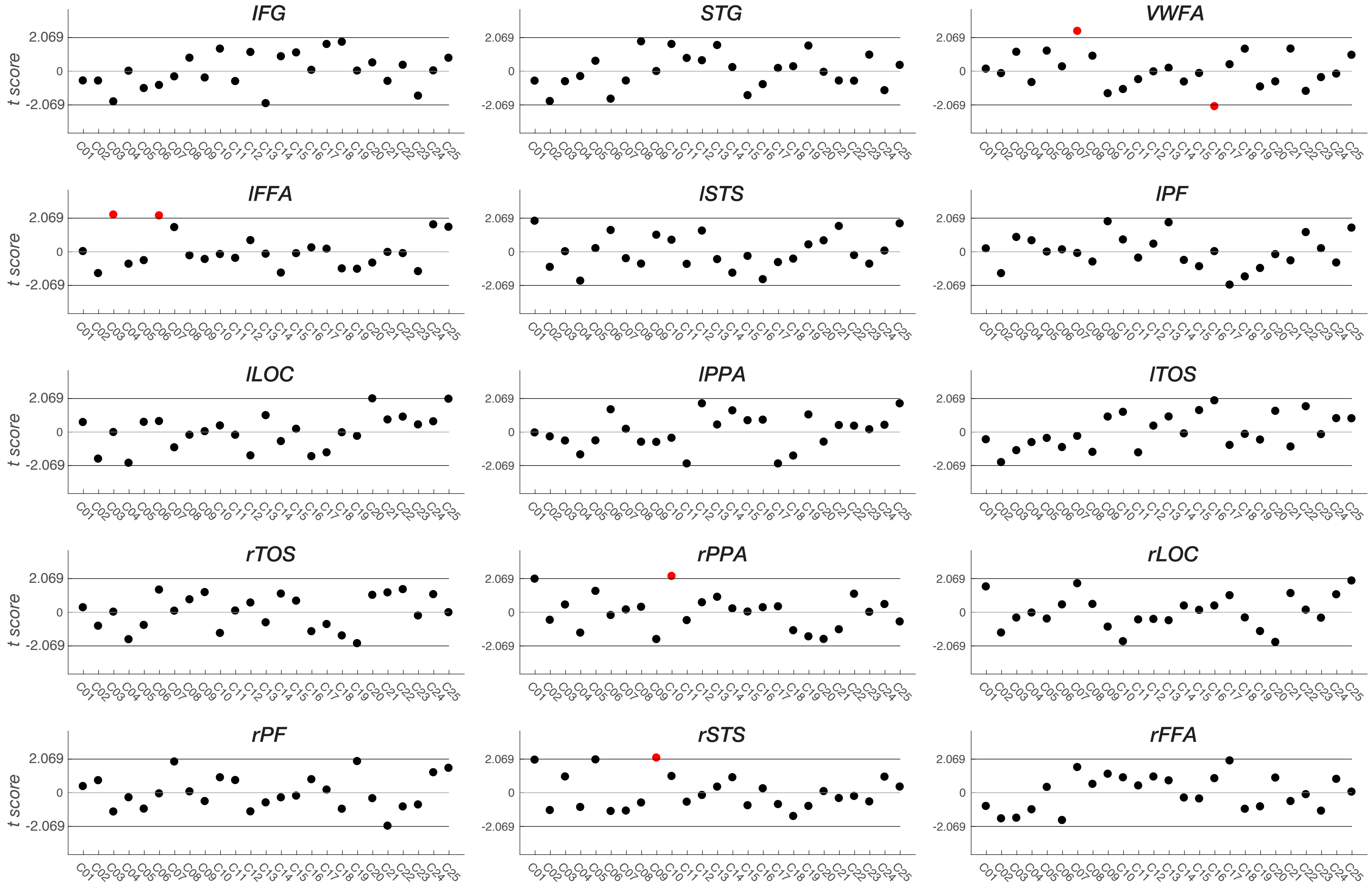


**Fig. S4. Crawford's t tests compared representational similarity of all category-selective ROIs in each control participant to the rest of the controls.** Red dots indicate significant deviations.

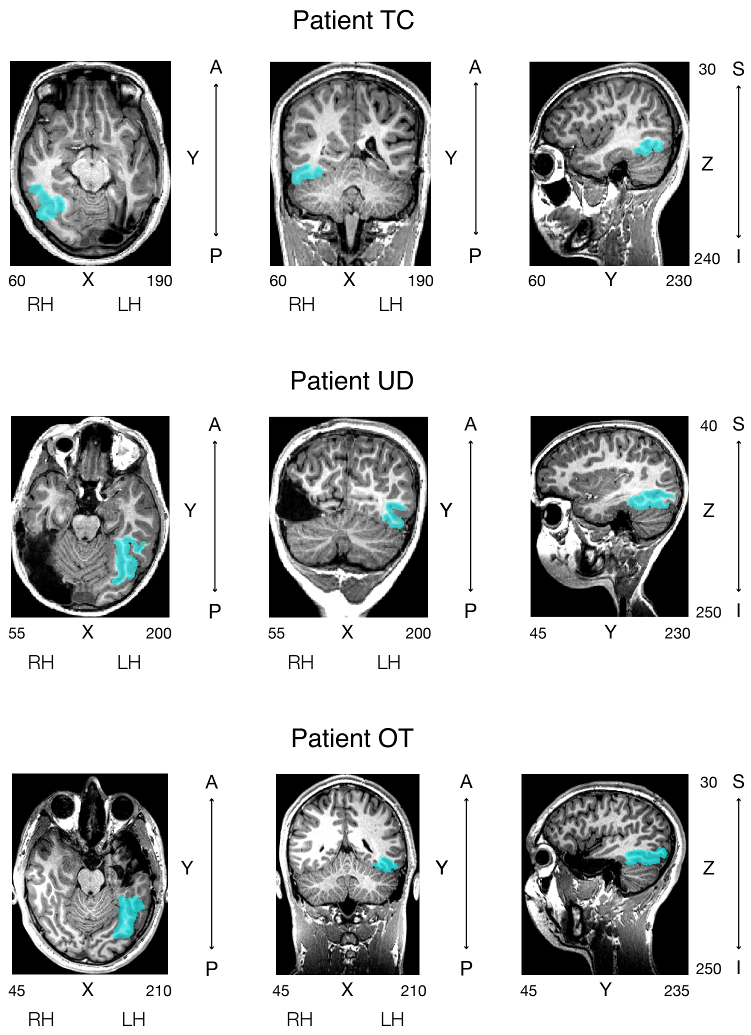


**Fig. S5. The anatomically defined right FG/OTS region visualized in corresponding volume space for TC, UD and OT.** Total number of anatomical voxels (1mm isotropic) is 7307 in TC, 12428 in UD, and 12013 in OT.
A = anterior, P = posterior, S = superior, I = inferior, RH = right hemisphere, LH = left hemisphere.

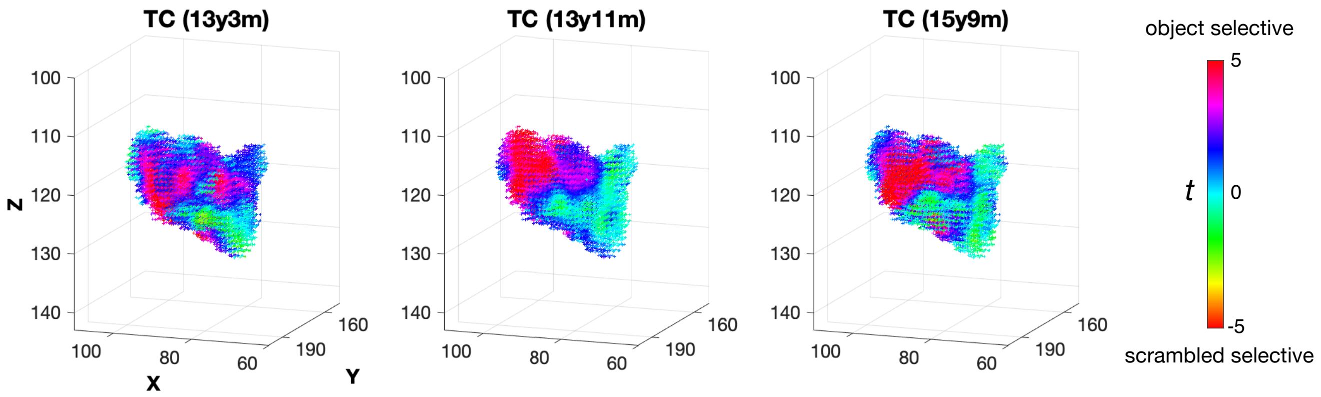


**Fig. S6. Longitudinal object versus scramble object selectivity in the anatomically defined FG/OTS in TC (3 sessions).** *t* score denotes changes over time in each voxel's selectivity of objects over scrambled objects. There were no significant differences between any two scan sessions, all $|t|$ values < 1.549, all $p$ values > 0.122, independent-samples $t$ tests at the voxel level.

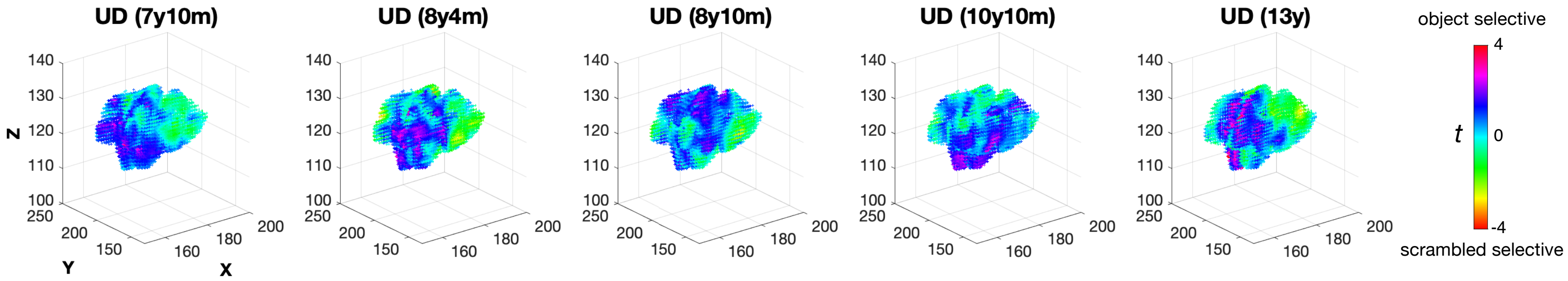


**Fig. S7. Longitudinal object versus scramble object selectivity in the anatomically defined FG/OTS in UD (5 sessions*).***
*t* score denotes changes over time in each voxel's selectivity of objects over scrambled objects. There were no significant differences between any two scan sessions (all $|t|$ values < 1.741, all $p$ values > 0.082), independent-samples $t$ tests at the voxel level.

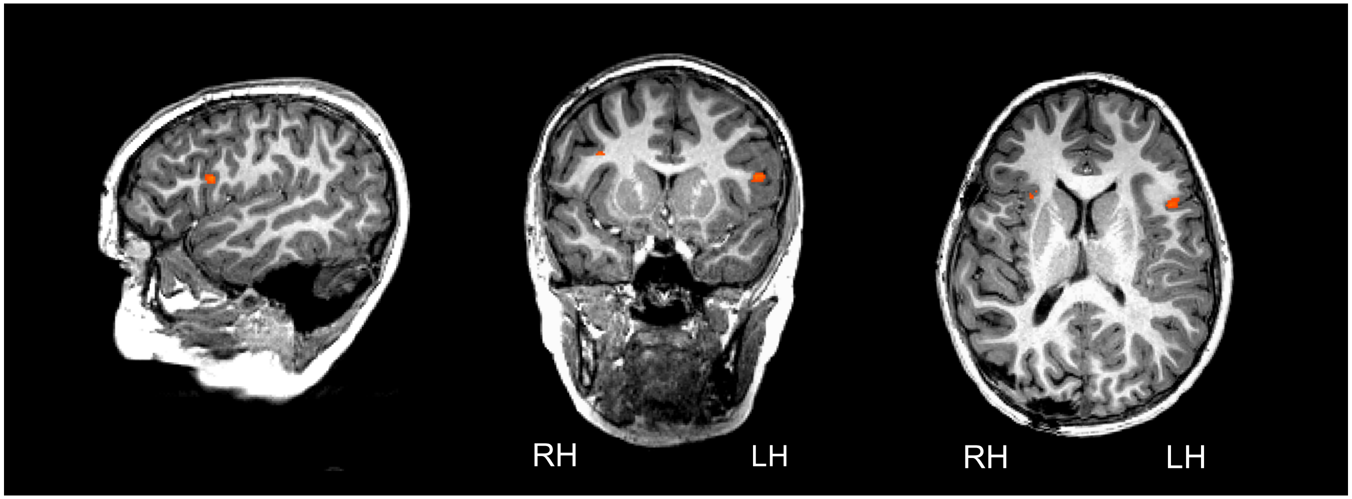


**Fig. S8. Post-surgical language mapping results in TC.**
Language-selective activation was found in the left Inferior Frontal Gyrus (Broca's area); no language-selective activation was found near the left Superior Temporal Gyrus (Wernicke's area) and no obvious language-selective activation was visible in the right hemisphere.

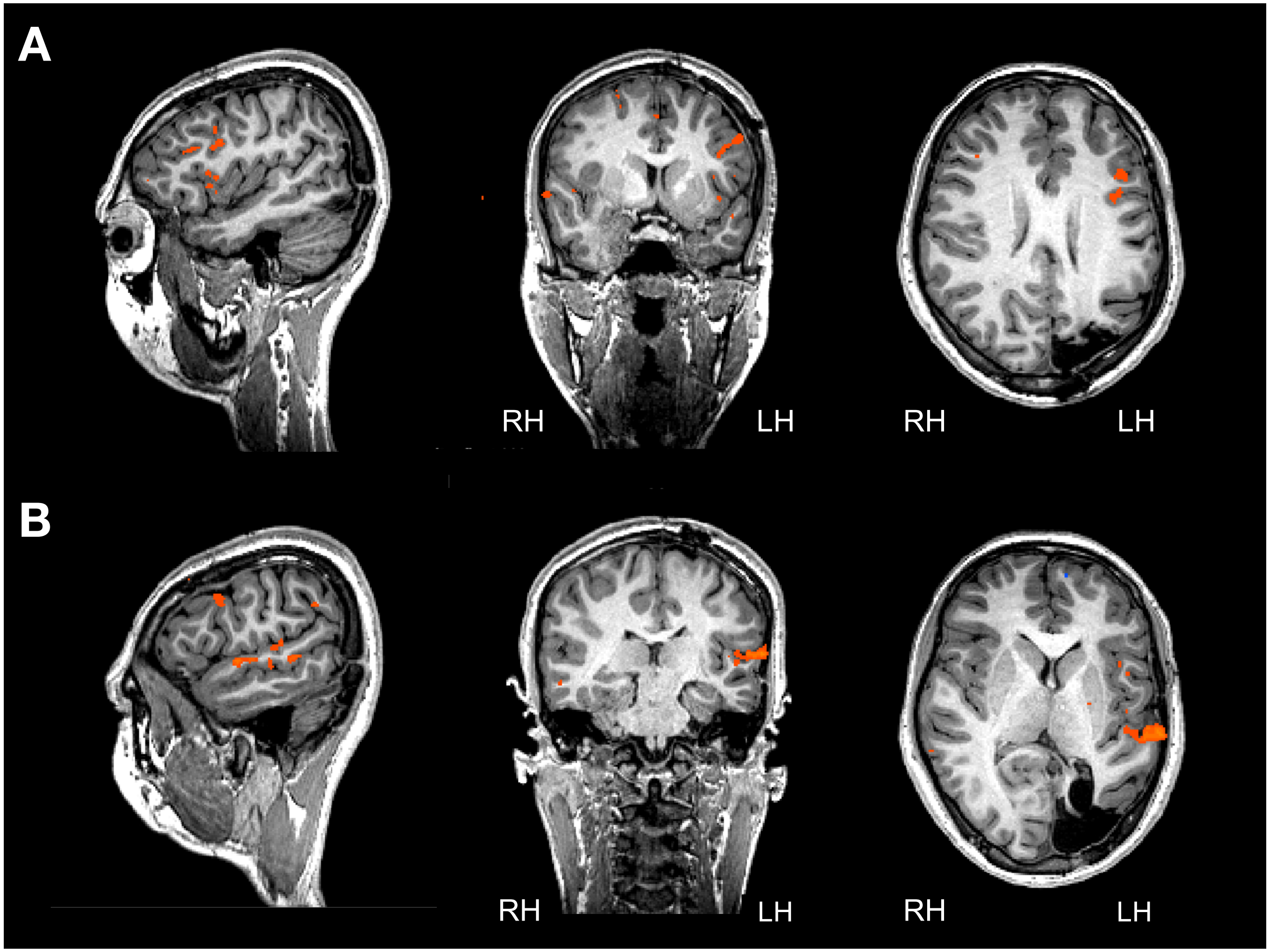


**Fig. S9. Post-surgical language mapping results in UD.**
A) language-selective activation in the left Inferior Frontal Gyrus (Broca's area)
B) language-selective activation in the left Superior Temporal Gyrus (Wernicke's area)

**Table S1. Demographic and surgical information for each of the five patients**

| Code | Gender | Diagnosis/Pathology | Preop MRI | Age at surgery | Lesioned hemisphere | Resection | Visual Fields | EESOS* |
|---|---|---|---|---|---|---|---|---|
| KN | M | N/A | N/A | 20m | left | left hemispherectomy | N/A | N/A |
| SN | M | N/A | N/A | 1d | left | evacuation of left temporal hematoma | N/A | N/A |
| TC | F | perinatal stroke with medically intractable focal epilepsy/multifocal encephalomalacia consistent with remote ischemic injury | abnormal MRI with lesion in the left occipital lobe | 13y3m | left | left posterior parietal and occipital lobectomy | right superior quadrantanopia | IID |
| UD | M | medically intractable focal epilepsy, symptomatic lesion/dysembryoplastic neuroepithelial tumor | abnormal MRI with lesion in the right temporal lobe | 6y9m | right | right medial inferior temporal lobe gross total tumor resection with posterior temporal and occipital lobectomy | left hemianopia | IA |
| OT | M | medically intractable focal epilepsy, symptomatic lesion/ganglioglioma, FCD | abnormal MRI with lesion in the medial left temporal lobe | 13y4m | left | medial left temporal lobe gross total tumor resection and left temporal lobectomy with preservation of medial structures | full | IA |

* EEESOS: Engle Epilepsy Surgery Outcome Scale

**Table S2. Patients' neuropsychological evaluation test performance pre- and post-surgery.**

| Code | Detailed IQ measures | Vision or visual motor integration | Memory & learning | Executive function | Academic skills/performance |
|---|---|---|---|---|---|
| KN | not done | not done | not done | not done | not done |
| SN | not done | not done | not done | not done | not done |
| TC | *Pre-surgery:*<br>WIAT III: $2^{nd}$ percentile*<br>*Post-surgery:*<br>not done | not done | not done | not done | PPVT: 1st percentile*<br>WIAT III:<br>Reading: 1st grade*<br>Spelling: 2nd grade* |
| UD | *Pre-surgery:*<br>WASI: 116 (full scale), 135 (verbal), 97 (performance)<br>*Post-surgery:*<br>WASI: 118 (full scale), 123 (verbal), 108 (performance) | Grooved pegboard:<br>$50^{th}$ percentile<br>(dominant hand) | not done | Working memory (from WISC-V):<br>*Post-surgery:*<br>$34^{th}$ percentile | WJ III ACH:<br>Reading: $63^{rd}$ percentile<br>Letter-Word: $67^{th}$ percentile<br>Passage: $56^{th}$ percentile<br>Calculation: $91^{st}$ percentile |
| OT | *Pre-surgery:*<br>WASI: 122 (full scale), 125 (verbal), 114 (performance)<br>*Post-surgery:*<br>WASI: 127 (full scale) | Grooved pegboard: average (dominant hand) | CVLT-C:<br>high average<br>WRAML-2:<br>high average | D-KEFS: superior | WJ III ACH: above age and grade expectancy |

* Could not be reliably obtained due to language barrier
PPVT: Peabody Picture Vocabulary Test
CVLT-C: California Verbal Learning Test–Children's Version
D-KEFS: The Delis–Kaplan Executive Function System
Grooved Pegboard: Grooved Pegboard for Manipulation and Dexterity Testing
WASI: Wechsler Abbreviated Scale of Intelligence
WIAT-III: Wechsler Individual Achievement Test–Third Edition
WISC-V: Wechsler Intelligence Scale for Children–Fifth Edition
WJ III ACH: The Woodcock-Johnson III Tests of Achievement
WRAML-2: Wide Range Assessment of Memory and Learning–Second Edition

**Table S3. Demographic information for each of the 25 fMRI controls.**

| Control | Gender | Age at scan | |
|---|---|---|---|
| | | year | month |
| C1 | F | 7 | 1 |
| C2 | M | 7 | 5 |
| C3 | M | 8 | 6 |
| C4 | F | 8 | 10 |
| C5 | M | 8 | 11 |
| C6 | F | 9 | 9 |
| C7 | M | 10 | 3 |
| C8 | M | 10 | 4 |
| C9 | F | 10 | 8 |
| C10 | M | 11 | 1 |
| C11 | M | 11 | 5 |
| C12 | M | 11 | 9 |
| C13 | M | 12 | 1 |
| C14 | F | 12 | 6 |
| C15 | M | 12 | 10 |
| C16 | F | 13 | 5 |
| C17 | M | 14 | 4 |
| C18 | F | 14 | 6 |
| C19 | F | 14 | 9 |
| C20 | F | 14 | 11 |
| C21 | M | 15 | 0 |
| C22 | F | 15 | 8 |
| C23 | M | 16 | 3 |
| C24 | M | 18 | 2 |
| C25 | M | 18 | 8 |
